# Physical Activity and Depressive Mood Shared the Structural Connectivity between Motor and Reward Network: a population-based study from the UK Biobank

**DOI:** 10.1101/2025.03.04.641424

**Authors:** Shiqi Di, Na Luo, Weiyang Shi, Zhengyi Yang, Jing Sui, Rongtao Jiang, Yue Cui, Zongchang Du, Jiaqi Zhang, Yawei Ma, Haiyan Wang, Congying Chu, Yuejia Zhong, Wen Li, Yuheng Lu, Hao Yan, Jinmin Liao, Dai Zhang, Vince Calhoun, Ming Song, Tianzi Jiang

**Author notes:** **Correspondence to: Tianzi Jiang,** Professor, Beijing Key Laboratory of Brainnetome and Brain-Computer Interface, Institute of Automation, Chinese Academy of Sciences, Beijing 100190, China. Shiqi Di and Na Luo contributed equally.

## Abstract

**Background:** Exercise has been revealed to leave a positive effect on alleviating depressive symptoms by various existing studies. However, the neural basis behind this phenomenon remains unknown, as well as its underlying biological mechanism.

**Aims:** Using a large neuroimaging cohort from UK Biobank, this study aimed to identify structural connectivity (SC) patterns simultaneously linked with physical activity and depression, as well as its biological interpretation.

**Method:** 492 participants with major depression disorder (MDD) and 535 healthy controls (HC) were extracted from the UK Biobank dataset. Partial least squares regression was first utilized to explore a replicable SC pattern simultaneously associated with physical activity and depressive mood. The neuromaps toolbox was applied to interpret the biological ontologies of the identified SC pattern. Neuroimaging-transcriptome association and enrichment analyses were conducted to explore its underlying genetic basis, pathways and cell types. The reproducibility and generalizability were tested on another independent MDD dataset (N=3,496) and bipolar disorder (BD) dataset (N=81).

**Results:** A SC pattern linked with exercise was identified to be both significantly correlated with depressive mood and group discriminative between MDDs and HCs, which primarily located between the motor-related regions and reward-related regions. This pattern was associated with multiple neurotransmitter receptors, such as serotonin and GABA receptors, and enriched in pathways like synaptic signaling and astrocytes cell type. The SC pattern and genetic results were also replicated in another independent MDD dataset and present commonalities with BD.

**Conclusions:** This study not only initially identified a reproducible shared SC pattern between physical activity and depressive mood, but also elucidated the underlying biological mechanisms, which enhanced our understanding of how exercise helps alleviate depression and may inform the development of novel neuromodulation targets.

## Introduction

Major depressive disorder (MDD) is a widespread mental disorder globally, characterized by persistent feelings of low mood and anhedonia. With a high incidence and recurrence rate, it substantially contributes to the global burden of disease^1^. Antidepressants are commonly prescribed for depression treatment, but challenges such as drug resistance and adverse reactions have been observed^2^. Consequently, there is an urgent need for novel and effective treatment strategies that could either substitute or complement conventional antidepressant medications.

Physical activity has been demonstrated as an effective non-pharmacological intervention for depression. For example, evidence from animal models demonstrates that exercise can induce antidepressant-like behavior, comparable to the effects of drug treatments^3,4^. Meta-analysis based on prospective studies have further revealed a reverse curve association between physical activity and the incidence of depression^5^, indicating potential preventive effects of exercise. Furthermore, a negative correlation has been observed between the frequency of physical activity and the severity of depression symptoms^6^.

However, the underlying neural basis linking exercise and depression have not been extensively studied on human brain. As the loss of pleasure in previous rewarding stimuli is a core symptom of MDD^7^, the function of reward-related regions and their association with mood have greatly aroused researchers’ attention^8^. Structural and functional alterations of brain regions responsible for reward processing, such as the prefrontal cortex, hippocampus, amygdala, basal ganglia, and thalamus, were well established in MDD^9–11^. These reward-related regions could also act as a predictor of treatment response in individuals with MDD^12,13^. Furthermore, the motor network has been shown to play a key role in antidepressant treatment response^14^. Although there are seldom studies directly exploring the macro-scale neuroimaging basis how exercise improves depressive mood, existed MRI studies suggested that physical activity could activate the function of motor-related regions^15,16^ and alter their connections with other brain regions^17,18^. Moreover, micro-scale experiments involving rotating wheel training on mice and rats indicated that exercise could increase the release of dopamine, serotonin and GABAergic neurotransmission in the reward-related brain regions^19–21^. These neurotransmitters play an important role in preventing the onset of depression or reducing depression^22^. Therefore, we hypothesized that the regulation in depressive mood resulting from physical activity may be associated with the connectivity between motor and reward networks.

Leveraging diffusion magnetic resonance imaging (dMRI) allows for the inference of structural connections (SC) within white matter, providing stable features of anatomical connections among diverse brain regions, which underpin the functional interactions across various brain regions^23^. Significant differences in SC have been identified in individuals with MDD^24^. In the context of physical activity, prior research has also found a connection between SC linking the frontal and temporal lobes and levels of physical activity^25,26^. Based on whole-brain SC, together with exercise assessment and depressive symptoms, our study seeks to elucidate potential neural mechanisms underpinning the association between physical activity and depressive mood.

In parallel, extensive studies have revealed depression-related alterations in neurotransmitters and their receptors, as well as dysregulation of astrocytes^27,28^, all of which have demonstrated responsiveness to exercise interventions^4,29,30^. These findings suggest that specific biological mechanisms might play a crucial role in linking exercise and depression. The Allen Human Brain Atlas (AHBA) dataset has been widely utilized for analyzing the transcriptomic basis associated with neuroimaging patterns^31^. Combining the gene expression dataset with neuroimaging patterns provides insights into understanding the microscale molecular basis and cell-specific information that underlie the macroscale imaging mechanisms^32,33^.

Collectively, we aim to conduct a multiscale analysis to systematically investigate how physical activity is linked to depressive status. We endeavor to elucidate the underlying brain patterns and biological factors that contribute to this association. The analysis pipeline was depicted in Figure 1. Leveraging dMRI data from a large sample of 1,027 participants (MDD/HC=492/535) obtained from the UK Biobank, we first employed a partial least squares (PLS) regression model to identify a SC pattern that exhibited both significant correlation with physical activity and group discriminative between MDDs and HCs. Subsequently, the neuromaps toolbox^34^ was employed to explore the neurotransmitter receptor profiles associated with the identified pattern. We then performed a neuroimaging-transcriptome association analysis to identify its underlying genetic basis, followed by pathway enrichment and cell type analysis. Finally, validation analyses were conducted on another independent MDD dataset and bipolar disorder (BD) dataset to explore the stability and generalizability of the findings.

**Figure 1.**
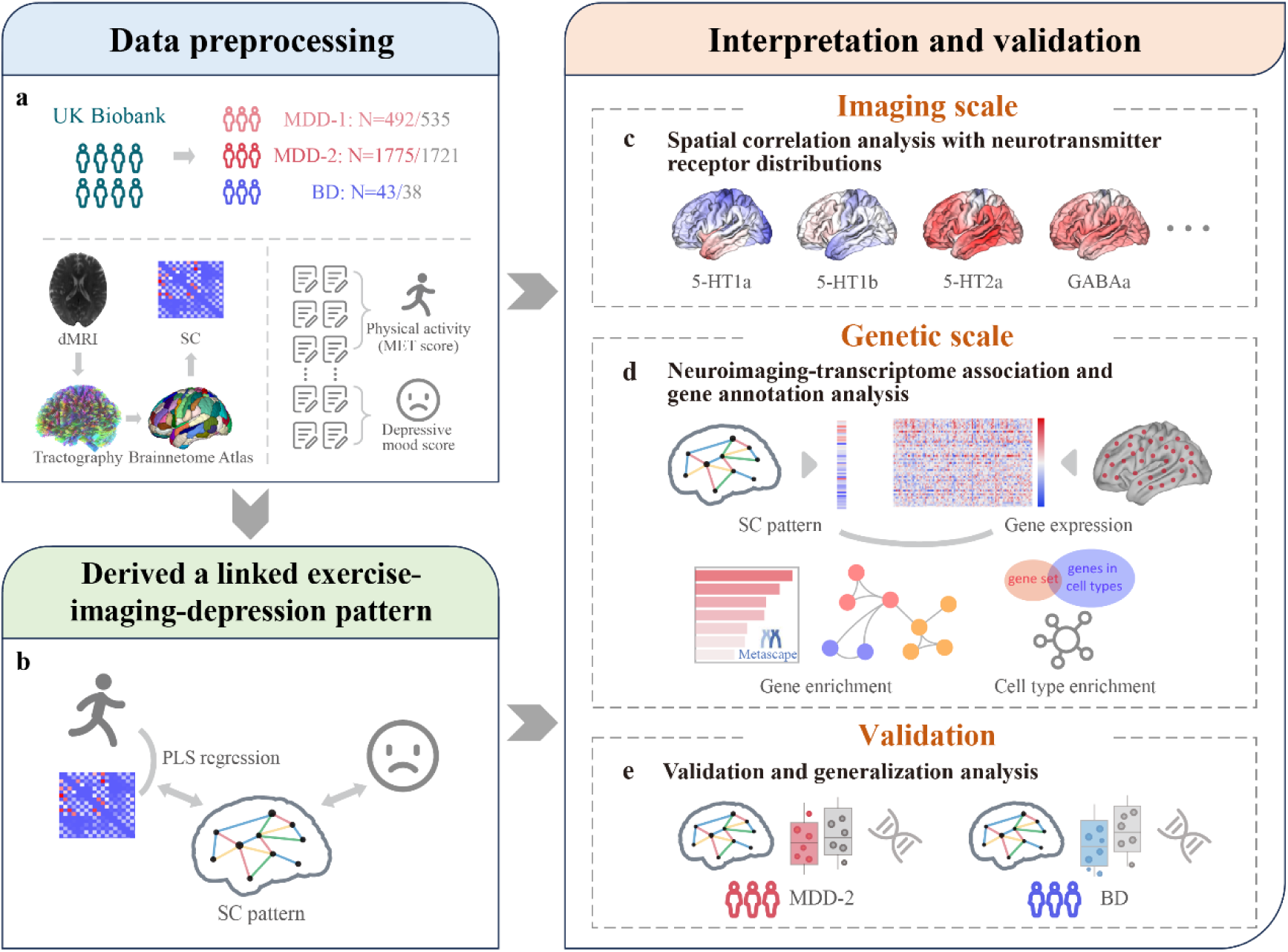
Overview of the analysis pipeline. **a**, Data preprocessing. MDD-1 dataset, MDD-2 dataset, and BD dataset were obtained from the UK Biobank. Based on dMRI data, probabilistic tractography was performed. Brainnetome Atlas (246 regions) was then aligned to extract the structural connectivity (SC) information at the brain region level. Alongside neuroimaging data, relative behavioral information (MET score and depressive mood score) was extracted from the responses to the touchscreen questionnaire. **b**, Derived a linked exercise-imaging-depression pattern. Based on MDD-1 dataset (MDD/HC=492/535), we performed PLS regression and identified a SC pattern that significantly associated with physical activity. Association analysis was also conducted to explore the relationship between this identified SC pattern and depressive mood. **c**, Spatial correlation analysis with neurotransmitter receptor distributions. The toolbox neuromaps was utilized to acquire neurotransmitter receptor maps, and subsequently, spatial correlations between the maps and the identified pattern were examined. **d**, Neuroimaging- transcriptome association and gene annotation analysis. We explored the genetic underpinnings of the identified SC pattern by integrating neuroimaging and transcriptome features. Gene enrichment and cell type enrichment analyses were then performed based on the identified gene list. **e**, Validation and generalization analysis. We conducted separate PLS regression analysis on another independent MDD-2 dataset and BD dataset. The SC patterns, group differences, correlations, and gene annotations were further analyzed on these datasets to evaluate the reproducibility and generalizability.

## Methods

### Participants

Three imaging datasets and one transcriptome dataset were included in the current study. The MDD-1 dataset, MDD-2 dataset and BD dataset were obtained from the UK Biobank (https://www.ukbiobank.ac.uk/). The UK Biobank study was approved by the North West Multi-centre Research Ethics Committee (https://www.ukbiobank.ac.uk/learn-more-about-uk-biobank/about-us/ethics) and all participants provided written informed consent. Inclusion criterions were listed below for each dataset.

#### (1) MDD-1 dataset

In the UK Biobank dataset, health-related outcomes included records of hospital inpatient obtained through external data linkage, coded according to the International Classification of Disease version 10 (ICD-10). From the entire cohort, 18,429 participants diagnosed with major depression disorder (MDD) (ICD-10: F32/F33) were selected. After excluding participants with missing relevant behavioral data and/or covariate variables, a total of 10,718 participants remained for association analysis, without considering MRI data.

We then constructed the MDD-1 dataset with individuals diagnosed with MDD described above and collected available diffusion imaging data. HCs were selected to match with the MDD participants based on age, sex, and site. HCs were defined as participants without any reported records of mental disorders or diseases of the nervous system according to ICD-10. After further exclusion of subjects who had missing relevant behavioral measures and/or covariates, and those with outlier data, the MDD-1 dataset finally included 1,027 subjects with 492 MDDs and 535 HCs. Detailed information regarding the data filtering process can be found in Figure 2.

**Figure 2.**
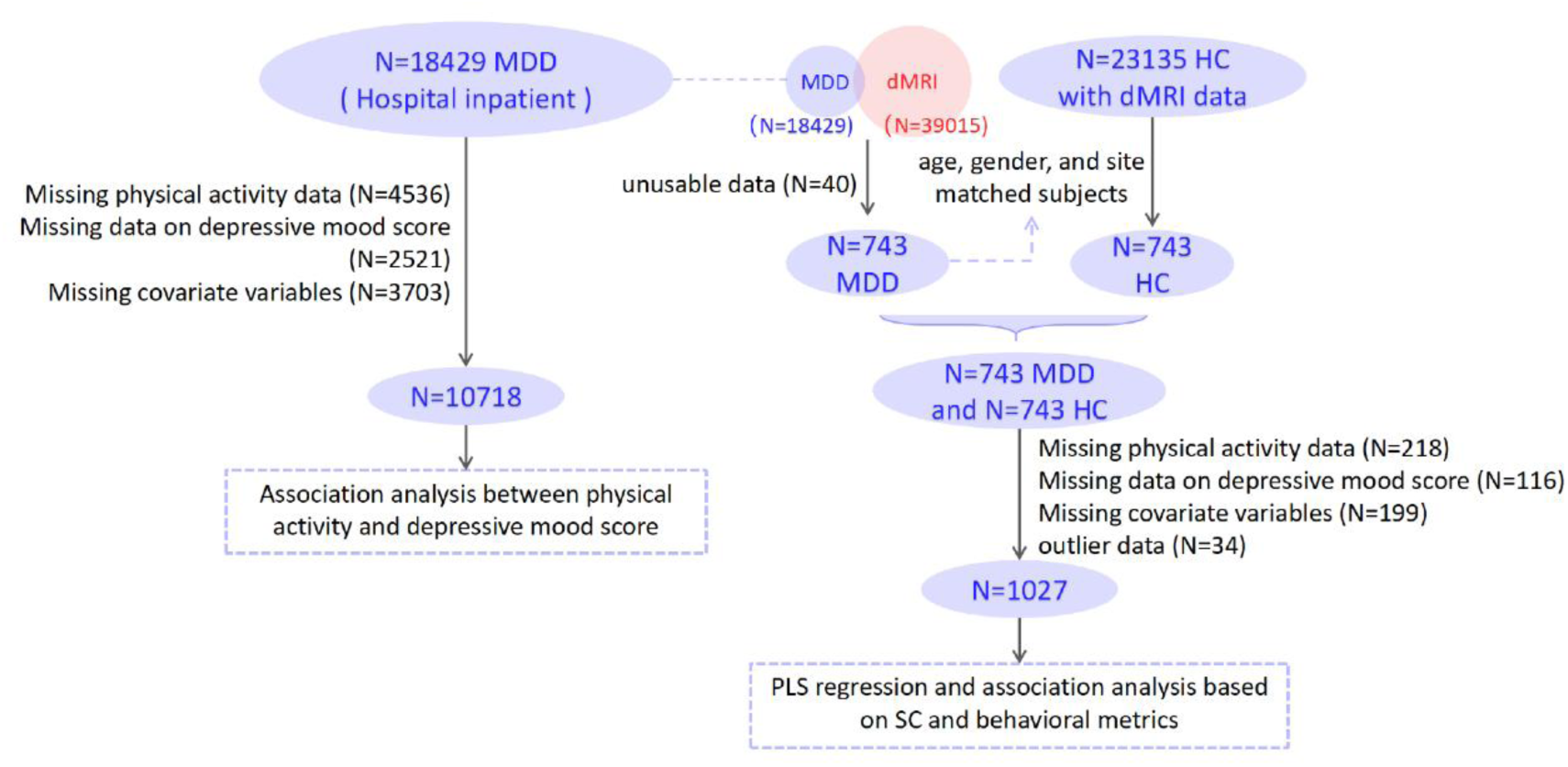
Flowchart of systematic sample selection for participants with MDD without considering MRI data (left) and for MDD-1 dataset (right).

#### (2) MDD-2 dataset

The selection criteria for participants with MDD in this dataset were determined based on the health outcomes characterized by Smith et al. (Field 20126)^35^. These outcomes were derived from a section of the touchscreen questionnaire and encompassed probable recurrent major depression (severe), probable recurrent major depression (moderate), and single probable major depression episode. In comparison to MDD-1 dataset, where individuals with depression were identified through hospital inpatient records, the diagnostic criteria employed in the MDD-2 dataset were less stringent, as it relied on responses obtained through the questionnaire. Consequently, MDD-2 dataset was employed for validation purposes.

To construct the MDD-2 dataset, individuals with depression were selected according to the criteria described above, and those with available dMRI data were retained. To ensure independence from the MDD-1 dataset, individuals already encompassed in the MDD-1 dataset were excluded from the MDD-2 dataset. After selection process, the MDD-2 dataset consisted of 1,775 MDDs and 1,721 HCs with corresponding age, sex, and site information.

#### (3) BD dataset

The BD dataset comprised participants who were diagnosed with BD according to their hospital records (ICD-10: F31). To ensure independence from the MDD-1 and MDD-2 dataset, individuals already encompassed in these two datasets were excluded from the BD dataset. Within this dataset, a total of 43 individuals with BD were included, and 38 HCs were selected, matched based on age, sex, and site.

#### (4) Transcriptome dataset

Gene expression data was from the Allen Human Brain Atlas (AHBA, https://human.brain-map.org) dataset^36^. The AHBA dataset included microarray data in 3,702 tissue samples of postmortem brains from 6 donors (five males and one female, aged 24-57 years old), and expression data for more than 20,000 genes were provided for each sample.

### Data collection and preprocessing

#### (1) Imaging dataset

The brain imaging data in UK Biobank were acquired using Siemens Skyra 3T scanners with the Siemens 32-channel head coil. T1-weighted structural imaging and dMRI data were utilized in this study. T1-weighted MRI data were obtained with a duration of 5 minutes, a resolution of 1×1×1 mm^3^ and a matrix of 208×256×256. The detailed scanning parameters for dMRI included a duration of 7 minutes, a resolution of 2×2×2 mm^3^ and a matrix of 104×104×72. The acquisition involved 100 distinct directions on two diffusion-weighted shells (b=1000, 2000 s/mm^2^), as well as 5 b=0 s/mm^2^ images.

The detailed imaging protocol and part of the processing pipeline for data from UK Biobank are provided in the online brain imaging document (https://biobank.ctsu.ox.ac.uk/crystal/ukb/docs/brain_mri.pdf) and elsewhere^37,38(p10)^. The preprocessing pipeline for dMRI data included correction of eddy currents and head motion using the FMRIB Software Library (FSL) (http://www.fmrib.ox.ac.uk/fsl/), followed by employing BEDPOSTx to estimate the orientation of fibers within each voxel^39,40^. To estimate connectivity probability between voxels, we performed probabilistic tractography using PROBTRACKx. Specifically, 5,000 streamlines were sampled for each voxel, and a threshold of 2/5,000 was applied to the output to reduce the occurrence of false positive samples^41^.

To establish SC at the brain region level, we aligned the Brainnetome Atlas^42^, which consists of 210 cortical and 36 subcortical regions, from MNI152 space to diffusion tensor imaging (DTI) space. Specifically, atlas was nonlinearly warped to the space of T1 using FNIRT (FMRIB’s Nonlinear Image Registration Tool) and subsequently transformed to the DTI space using Advanced Normalization Tools (ANTS)^43^. We calculated the average SC between brain regions based on the Brainnetome Atlas, resulting in a symmetric SC matrix of 246×246 for each individual. Extracting the upper triangular elements of the matrix, we obtained 30,135 edges that served as SC features for subsequent analysis.

#### (2) Physical activity measurement

Metabolic Equivalent Task (MET) score was used to measure physical activity level, based on the International Physical Activity Questionnaire (IPAQ) guidelines^44^. According to the protocol for IPAQ short form, total physical activity included three types of activities: walking, moderate activity, and vigorous activity. The MET score for each type was calculated by multiplying the duration (minutes/day) by the number of days on a typical week. The total physical activity (MET-minutes/week) was obtained by combining the three types of activities, where a value of 3.3 METs was assigned to walking, 4 METs to moderate activity, and 8 METs to vigorous activity^45^. In addition, data with a total duration exceeding 960 minutes across the three categories were treated as outliers, and any record with a duration less than 10 minutes was set to 0. The IPAQ truncation rules specified that all variables with a duration above 180 minutes in any of the three activity categories were truncated and replaced with 180 minutes in a new variable.

#### (3) Depression measurement

The depressive mood score was assessed based on participants’ responses to touchscreen questionnaire about frequency of depressed mood, disinterest, tenseness and tiredness in last 2 weeks^46^. Participants’ responses included: “Not at all,” “Several days,” “More than half the days,” and “Nearly every day,” which were coded from 0 to 3 according to the Patient Health Questionnaire (PHQ-9)^47^. The sum of the responses from four questions was used to calculate the depressive mood score that ranged from 0 to 12, reflecting the severity of the depressive mood.

#### (4) Transcriptome dataset

The microarray data were preprocessed utilizing the toolbox abagen^48^ and spatially matched to 246 brain regions as defined in the Brainnetome Atlas. Since the absence of tissue samples in 11 out of the 246 brain regions, a gene expression matrix of 235 × 15,605 was obtained, where 235 refers to the number of brain regions and 15,605 to the number of genes in each region. The details of preprocessing pipeline are provided in supplemental material.

### Association between physical activity and depressive mood score

A general linear model (GLM) was utilized to examine the relationship between physical activity and depressive mood score among a group of 10,718 participants with MDD without considering MRI data. In this analysis, the depressive mood score served as the dependent variable, while physical activity and several covariate variables were modeled as fixed effects. The covariate variables encompassed several factors such as age, sex, educational attainment (college or university degree/other), ethnicity (white/non-white), Townsend deprivation index, employment status (in paid employment/other), body mass index, and the average total household income before tax (<£18,000, £18,000-£30,999, £31,000-£51,999, £52,000-£100,000, >£100,000; encoded using four-column dummy variable). All behavioral variables and covariates were obtained at the baseline, corresponding to the initial assessment visit.

### Revealing neuroimaging patterns underlying the association between physical activity and depressive mood score

We used PLS regression to assess the association between physical activity and whole-brain structural connections, with the total MET scores serving as the response variables and the inter-regional SC features as the predictor variables^49,50^. A permutation test was repeated 1,000 times to test the statistical significance of the components’ explained variance. In addition, bootstrap resampling was conducted 1,000 times to assess the contribution of each SC to the components. The contribution was represented using Z scores, calculated by dividing the PLS weight of each connectivity by its bootstrapped standard error.

The SC latent component (sc-LC) derived from PLS which achieved statistical significance and explained the highest variance was retained for the subsequent analyses. The sc-LC score was calculated by multiplying the whole-brain structural connections by the weights of sc-LC. The group difference between MDDs and HCs of the sc-LC score was tested using two-sample t-test. We then calculated the correlation between the sc-LC score and the depressive mood score. The covariates in the correlation analyses included group information, age, sex, assessment center for imaging, educational attainment, ethnicity, Townsend deprivation index, employment status, body mass index, and the average total household income before tax. The detailed fields corresponding to the variables were presented in Table S1. The top contributing SC with |Z| > 3 were further selected to interpret the imaging variations associated with physical activity^32,51^.

### Spatial correlation analysis with neurotransmitter receptor distributions

We employed the neuromaps toolbox^34^ to detect the neurotransmitter receptors associated with the SC pattern. To accomplish this, we calculated the node strength at the regional level based on |Z| of each SC within the sc-LC and regarded the spatial distribution of the node strength as the SC pattern. The neurotransmitter receptor maps were obtained through the neuromaps toolbox^52,34^, with each map sourced from previous positron emission tomography (PET) studies. Details for each receptor map can be found in supplemental material and Table S2. These neurotransmitter receptor maps included maps of serotonin receptors (5-HT1a, 5-HT1b, 5-HT2a, 5-HT4, 5-HT6), dopamine receptors (D1, D2), acetylcholine receptors (α4β2, M1), GABA receptor (GABAa), glutamate receptor (mGluR5), cannabinoid receptor (CB1), mu-opioid receptor (MOR), and histamine receptor (H3). We employed a weighted average to combine maps obtained from different studies that corresponded to the same neurotransmitter receptor^52^ and subsequently mapped them to the Brainnetome Atlas. Pearson correlation was then applied to determine the statistical association between the SC pattern and each of these receptor maps. In order to assess the significance of these associations, spatial null models (spin test) were employed^53,54^. Additionally, p-values were corrected for multiple comparisons using the FDR method.

### Neuroimaging-transcriptome association analysis

To further assess the genetic basis associated with the discovered sc-LC pattern, we used the prepared gene expression matrix (235 regions × 15,605 genes) obtained from the AHBA. We separated the positive and negative Z scores of the sc-LC and calculated the node strength of Z scores at the regional level for positive and negative condition, respectively. We then employed PLS regression to dig out the gene expression pattern that associated with the sc- LC pattern. The gene expression data, covering 235 brain regions, served as predictor variables, and the node strength of the sc-LC corresponding to these 235 ROIs were retained as the response variables. We performed a permutation test (n=1,000) to assess the significance of the variance explained by the PLS component. After selecting the genetic components (g-LC) that exhibited statistical significance and the largest explained variance, we conducted bootstrapping to calculate the Z score, the ratio of the weight to the bootstrapped standard error, for each gene. Finally, genes with |*Z*| > 3 were retained for subsequent gene enrichment analysis^32,51^.

### Genetic pathway and cell type analysis

We adopted the Metascape toolbox^55^ to perform Gene Ontology (GO) analysis, Kyoto Encyclopedia of Genes and Genomes (KEGG) pathway and DisGeNET enrichment analysis. All genes in the genome were selected as the reference gene list. The significance values were FDR-adjusted to correct for multiple comparisons.

Furthermore, a cell-type enrichment analysis was conducted to identify the specific cell types that were enriched for these genes. Cell types included seven canonical cell classes: astrocytes, endothelial, microglia, excitatory neurons, inhibitory neurons, oligodendrocytes, and oligodendrocyte precursor cells, as reported in the study by Seidlitz et al.^56^. To calculate the ratio of overlap between each cell type’s gene set and our gene list, we divided the number of overlapped genes by the total number of genes included in our gene list for each cell type. A permutation test was performed to calculate the p-value corresponding to the ratio for each cell type. All the p-values are corrected using the FDR.

## Results

### A SC pattern associated with physical activity and depressive mood

Leveraging a large cohort from the UK Biobank, we employed the general linear model (GLM) to examine the association between physical activity and depressive mood (details in supplemental material). Individuals with a documented history of depression in the records of hospital inpatient were chosen to enhance the rigor of subject selection. Based on the baseline scores of 10,718 depressed subjects without considering MRI data, we found that increased weekly physical activity was linearly linked to improved depressive mood after adjusting for relevant covariates (*t* = -8.65, *p* = 6.0e-18) (Figure 3a). Considering the depressed individuals with available dMRI data, along with age-, sex-, and site-matched HCs, we constructed the MDD-1 dataset (N=1,027, MDD/HC=492/535) for subsequent primary analyses. The process of data filtering is illustrated in Figure 2. The demographic characteristics of the participants in MDD-1 dataset are presented in Table 1. The association above remained (*t* = -5.76, *p* = 1.1e-8) in MDD-1 dataset. The physical activity score (Cohen’s *d* = -0.19, *p* = 3.1e-3) and depressive mood score (Cohen’s *d* = 0.69, *p* = 2.7e-25) exhibited significant differences between MDDs and HCs (Figure S1).

**Figure 3.**
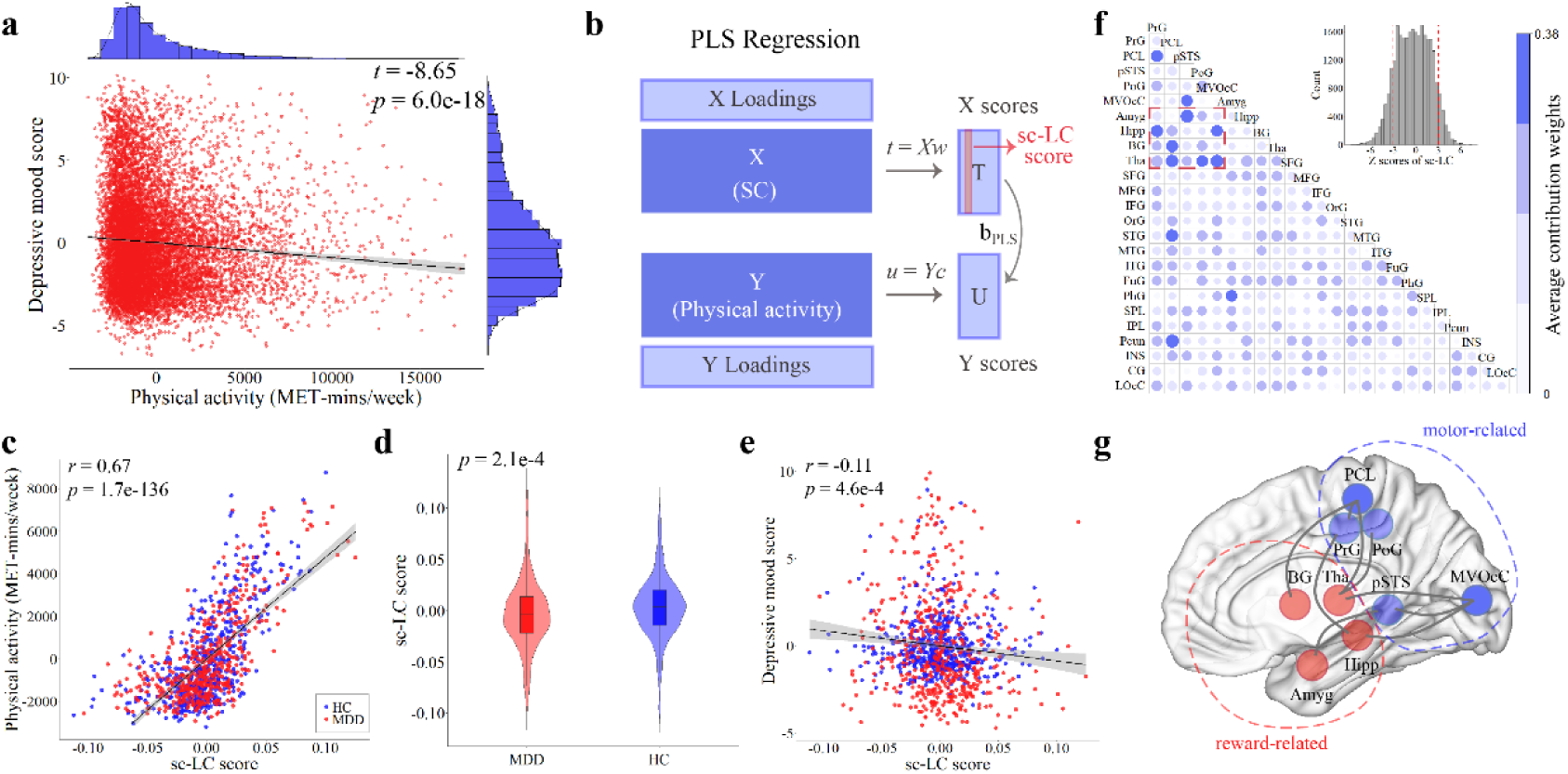
A linked exercise-imaging-depression pattern identified using PLS regression. **a**, Association analysis between physical activity and depressive mood score (N=10,718). The error band indicates a 95% confidence interval. **b**, Schematic overview of PLS Regression. Structural connections were used as predictor variables and physical activity (MET score) as response variables. The vectors *w* and *c* captured the weights of latent component for X and Y, and matrices T and U contained the component scores of X and Y respectively. The sc-LC, which explained the largest percentage of variance in Y, was utilized. **c**, Correlation between the sc-LC score (a weighted linear combination of 30,135 whole-brain structural connections) and physical activity (MET score) based on the MDD-1 dataset (MDD/HC=492/535). The red dots in the scatter plot indicate subjects with MDD, and the blue dots represent HC. The error band indicates a 95% confidence interval. **d**, Two-sample two-tailed t-test of sc-LC score between the MDDs and the HCs. Violin and box plots were used to show the distribution of sc-LC scores for the two groups. In each box plot, the midline indicated the median, the box was bounded by the first and third quartiles, and the whiskers extended to 1.5 × interquartile range. **e**, Correlation between the sc-LC score and depressive mood score. **f**, Average contribution weights calculated based on reliable connections (upper right) retained by Z scores of the sc-LC. 246 subregions were grouped into 24 macroscale brain regions based on the anatomical definition of the Brainnetome Atlas, and the contribution weight was defined as the number of connections with reliable contributions (|Z| > 3) divided by the total number of connections between two macroscale regions. Larger and darker dots represent higher weights. **g**, Connections with high weights in **f** were mainly concentrated between reward-related and motor-related regions. PrG, precentral gyrus; PCL, paracentral lobule; pSTS, posterior superior temporal sulcus; PoG, postcentral gyrus; MVOcC, medioventral occipital cortex; Amyg, amygdala; Hipp, hippocampus; BG, basal ganglia; Tha, thalamus; SFG, superior frontal gyrus; MFG, middle frontal gyrus; IFG, inferior frontal gyrus; OrG, orbital gyrus; STG, superior temporal gyrus; MTG, middle temporal gyrus; ITG, inferior temporal gyrus; FuG, fusiform gyrus; PhG, parahippocampal gyrus; SPL, superior parietal lobule; IPL, inferior parietal lobule; Pcun, precuneus; INS, insular gyrus; CG, cingulate gyrus; LOcC, lateral occipital cortex.

**Table 1.** Demographic Characteristics of Participants.

|  | MDD-1 Dataset |  |  | MDD-2 Dataset |  |  | BD Dataset |  |  |
| --- | --- | --- | --- | --- | --- | --- | --- | --- | --- |
|  | MDD | HC | p | MDD | HC | p | BD | HC | p |
| **Subjects** | **492** | **535** |  | **1775** | **1721** |  | **43** | **38** |  |
| **Age, Years** | 62.10<br>±<br>7.66 | 62.07<br>±<br>7.53 | 0.95 | 62.50<br>±<br>7.38 | 62.24<br>±<br>7.31 | 0.30 | 62.61<br>±<br>7.44 | 62.76<br>±<br>7.40 | 0.93 |
| **Sex (Male/Female)** | 39%/<br>61% | 36%/<br>64% | 0.43 | 40%/<br>60% | 40%/<br>60% | 0.79 | 47%/<br>53% | 50%/<br>50% | 0.75 |
| **Education attainment (High/Low)** | 47%/<br>53% | 58%/<br>42% | <0.001 | 54%/<br>46% | 54%/<br>46% | 0.94 | 56%/<br>44% | 63%/<br>37% | 0.50 |
| **Ethnicity (White/Non-white)** | 98%/<br>2% | 98%/<br>2% | 0.36 | 98%/<br>2% | 97%/<br>3% | 0.27 | 98%/<br>2% | 100%/<br>0% | 0.34 |
| **Townsend deprivation index** | -1.36<br>±<br>2.99 | -2.14<br>±<br>2.63 | <0.001 | -1.69<br>±<br>2.66 | -2.05<br>±<br>2.67 | <0.001 | -0.69<br>±<br>3.08 | -2.63<br>±<br>1.61 | <0.001 |
| **Employment status (In paid employment/Other)** | 39%/<br>61% | 48%/<br>52% | 0.003 | 45%/<br>55% | 48%/<br>52% | 0.14 | 28%/<br>72% | 50%/<br>50% | 0.04 |
| **Body mass index** | 28.44<br>±<br>5.22 | 25.77<br>±<br>4.16 | <0.001 | 26.87<br>±<br>4.37 | 26.25<br>±<br>4.39 | <0.001 | 28.32<br>±<br>4.62 | 26.55<br>±<br>3.49 | 0.06 |
| **Average total household income before tax** |  |  |  |  |  |  |  |  |  |
| **Less than 18,000** | 24% | 9.5% | <0.001 | 12% | 8% | <0.001 | 28% | 13% | 0.20 |
| **18,000 to 30,999** | 32% | 24% |  | 28% | 25% |  | 26% | 24% |  |
| **31,000 to 51,999** | 26% | 29% |  | 31% | 31% |  | 32% | 32% |  |
| **52,000 to 100,000** | 15% | 28% |  | 23% | 26% |  | 14% | 26% |  |
| **Greater than 100,000** | 3% | 9.5% |  | 6% | 10% |  | 0% | 5% |  |
| **Physical activity, MET-min/week** | 2207<br>±<br>2177 | 2791<br>±<br>2195 | <0.001 | 2622<br>±<br>2204 | 2774<br>±<br>2239 | 0.04 | 1981<br>±<br>1712 | 2800<br>±<br>1732 | 0.04 |
| **Depressive mood score** | 3.35<br>±<br>3.27 | 1.03<br>±<br>1.54 | <0.001 | 1.64<br>±<br>1.82 | 0.94<br>±<br>1.37 | <0.001 | 2.60<br>±<br>2.66 | 0.89<br>±<br>1.23 | <0.001 |

Note: Values are presented as mean ± SD for continuous variables and as percentage for categorical variables. P values were obtained using two-sample two-tailed t-test for continuous variables and Chi-square test for categorical variables.

Using PLS regression (Figure 3b), we identified a SC latent component (sc-LC) that was associated with physical activity and explained the largest percentage of variance (44%, permuted *p* = 1.0e-3) in MET score. After regressing group differences, the sc-LC score, which was calculated by multiplying the whole-brain structural connections by the weights of sc-LC, still exhibited a significant correlation with MET score (*r* = 0.67, *p* = 1.7e-136) (Figure 3c). The MDD group showed significantly lower sc-LC score than the HC group (Cohen’s *d* = -0.23, *p* = 2.1e-4) (Figure 3d). These findings suggest that increased weekly exercise time is associated with higher sc-LC score, which tends to represent healthier states. Using different intensities of physical activity (walking, moderate activity, and vigorous activity), we found that all types of physical activity significantly correlated with the sc-LC score, with moderate activity showing the strongest correlation (see supplemental material and Figure S2 for more details). Furthermore, we found that the sc-LC score was negatively correlated with depressive mood score (*r* = -0.11, *p* = 4.6e-4) (Figure 3e). Given that the sc- LC score was simultaneously associated with the MET score and the depressive mood score, we performed a mediation analysis^57^ to explore whether a mediation effect of *MET* 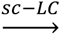 *mood* may occur. However, the indirect effect (CI = [-0.01, 0.09]) was not significant (for details refer to the supplemental material).

To better interpret and visualize the sc-LC pattern, we grouped the 246 brain regions into 24 larger regions according to the anatomical definition of the Brainnetome Atlas. We found that the structural connections with significant contributions were primarily located between the reward circuit where amygdala, hippocampus, basal ganglia and thalamus are located^58,59^, and regions involved in motor such as precentral gyrus, paracentral lobule, postcentral gyrus, posterior superior temporal sulcus, and medioventral occipital cortex^60,61^ (Figure 3f, g).

To examine the potential influence of inter-group differences in physical activity on latent structural connectivity when combining MDD and HC, we conducted PLS analysis on the MDD/HC group separately. The identified patterns from both populations presented high similarity with the original pattern (*r* = 0.69, *p* < 0.001 for the MDD group; *r* = 0.67, *p* < 0.001 for the HC group), replicating the connections between the motor and reward networks (see supplemental material and Figure S3 for more details). These results indicated that the group difference in physical activity between MDD and HC has little or at best a very subtle effect on the identified latent structural connectivity pattern. Moreover, to assess the potential medication effects on our results, we followed Glanville et al.^62^ to divide the MDD-1 dataset into a medicated group and an unmedicated group (Table S3). PLS analyses were conducted separately on the medicated group and unmedicated group. As shown in Figure S4, the derived sc-LC pattern of both sub-groups replicated the connection between motor and reward network as our primary findings using the whole MDD-1 dataset, suggesting that the influence of medication treatment on the outcomes was relatively minor (for details refer to the Supplementary Results).

### Neurotransmitter receptor maps associated with the SC pattern

We used the neuromaps toolbox^34^ to download the neurotransmitter receptor maps, which were sourced from prior positron emission tomography (PET) studies. For the obtained neurotransmitter receptor maps, significant spatial correlations were observed between the identified sc-LC pattern and serotonin receptor 5-HT1a (*r* = -0.30, false discovery rate (FDR)- corrected *p* < 0.01), 5-HT2a (*r* = -0.33, FDR-corrected *p* < 0.01), acetylcholine receptor α4β2 (*r* = 0.23, FDR-corrected *p* < 0.05), M1 (*r* = -0.21, FDR-corrected *p* < 0.05), GABA receptor GABAa (*r* = -0.32, FDR-corrected *p* < 0.01), and glutamate receptor mGluR5 (*r* = -0.25, FDR-corrected *p* < 0.05) (Figure 4a).

**Figure 4.**
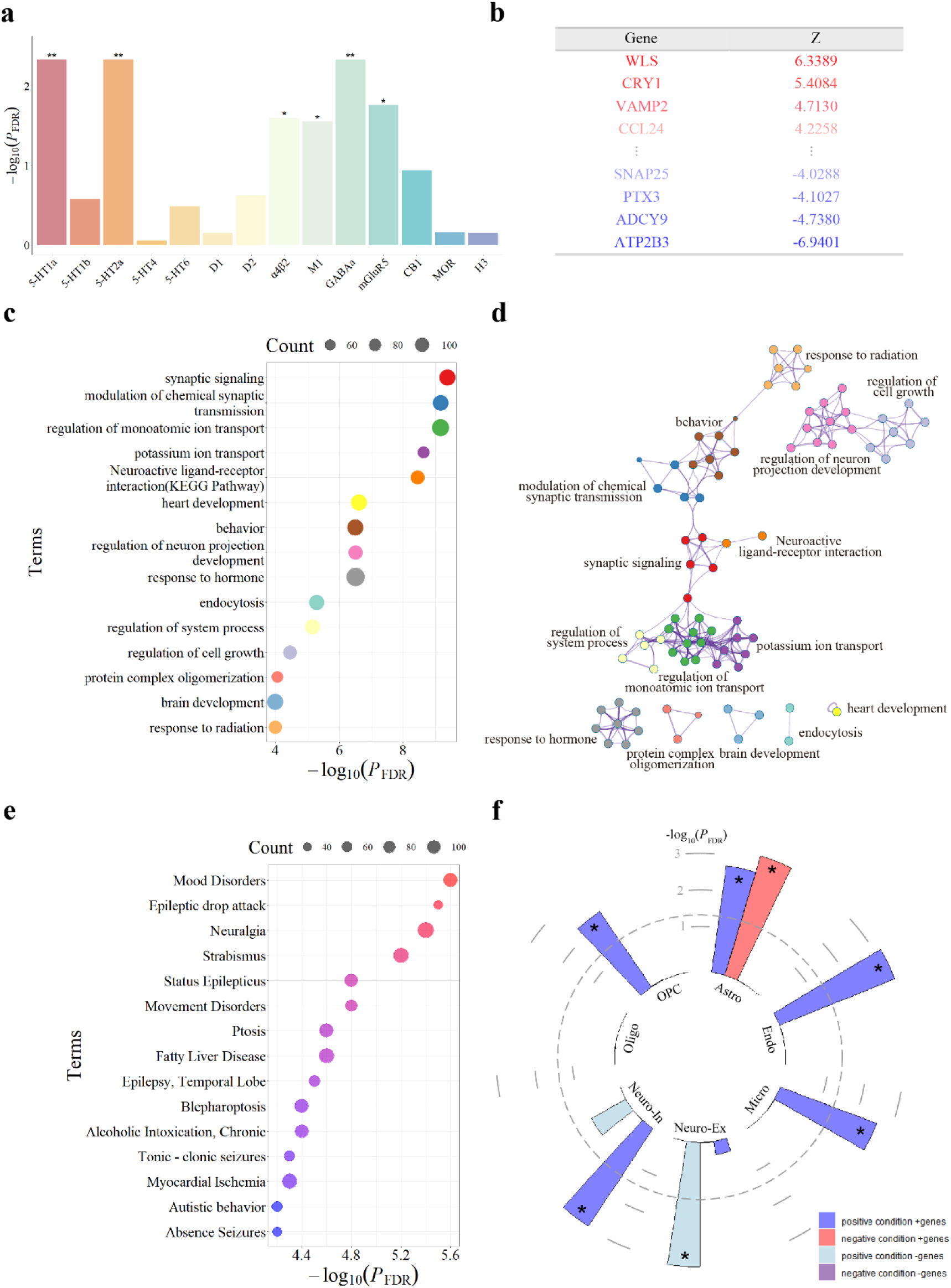
Spatial correlation with neurotransmitter receptors and neuroimaging- transcriptome association analysis to reveal the biological basis of the pattern. **a**, Spatial correlation analysis between the sc-LC pattern and neurotransmitter receptor maps. The statistical significances of correlations were assessed using spatial null models^53,54^, corrected for multiple comparisons using the FDR. The asterisk denoted the correlation was significant (*FDR-corrected *p* < 0.05, **FDR-corrected *p* < 0.01). **b**, Gene list for g-LC with genetic weights |Z| >3 for subsequent enrichment analysis. **c**, The top 15 representative enriched terms of GO biological processes and KEGG pathway for g-LC gene list. The size of each dot corresponds to the number of genes involved in the given ontology terms. **d**, Network visualization of enriched terms. Each node in the network corresponds to an enriched term, with nodes belonging to the same cluster being color-coded identically. **e**, The top 15 representative terms enriched in DisGeNET for g-LC gene list. The size of each circle corresponds to the number of genes involved in the given ontology terms. **f**, Enrichment analysis for cell types. Positive and negative conditions corresponded to positive and negative Z scores for sc-LC, respectively. In specific condition, genes exhibiting a positive correlation with the node strength of Z scores were referred to as “+genes”, whereas those demonstrating a negative correlation were termed “-genes”. P value was corrected by FDR. Astro, astrocytes; Endo, endothelial; Micro, microglia; Neuro-Ex, excitatory neurons; Neuro-In, inhibitory neurons; Oligo, oligodendrocytes; OPC, oligodendrocyte precursor cells.

### Gene expression profiles underlying the SC pattern

Based on the preprocessed transcriptome data from AHBA dataset^36^ (235 regions × 15,605 genes), we used PLS regression to identify the gene expression pattern associated with the sc-LC pattern. The sc-LC pattern was characterized by both positive and negative conditions, with the positive condition determined by the node strength of positive Z scores of sc-LC, and the negative condition defined by the node strength of negative Z scores. The explained variances and statistical significance of all PLS-derived genetic latent components (g-LC) for each condition are listed in Table S4. Two significant g-LC patterns were identified in the positive condition and one significant g-LC pattern was observed in the negative condition.

We adopted the g-LC patterns that explained the largest proportion of variance in the node strength of Z scores of sc-LC (16%, permuted *p* = 4.4e-2 for positive condition; 17%, permuted *p* = 2.0e-3 for negative condition) and showed the highest correlation with the sc- LC pattern for both positive and negative conditions (*r* = 0.40, *p* = 2.4e-10 for positive condition; *r* = 0.42, *p* = 2.6e-11 for negative condition) as the main patterns for the subsequent analysis (Figure S5a, c). Based on the Z scores calculated from g-LC using bootstrapping, 1,487 genes with |Z| > 3 for positive condition and 1,394 genes with |Z| > 3 for negative condition were extracted for further analysis (Figure S5b, d). Validation analyses considering the effects of potential sampling bias in donors of the AHBA dataset further suggested that the identified genetic components were stable to hemisphere and sex (for details refer to the supplemental material). In addition, the second significant g-LC pattern for the positive condition explained 13% of the variance (permuted *p* = 4.3e-2) and was significantly correlated with the sc-LC pattern (*r* = 0.36, *p* = 2.0e-8) (Figure S6a, b). For this component, 1,317 genes with |Z| > 3 were extracted for subsequent analysis.

### Genetic pathway and cell type analysis

The gene list (Figure 4b) extracted from the main g-LC patterns was employed for enrichment analysis with Metascape, focusing on GO biological processes, KEGG pathway, and DisGeNET category. Enrichment analysis revealed the top 15 significant terms, including GO biological processes “synaptic signaling”, “modulation of chemical synaptic transmission”, “regulation of monoatomic ion transport”, “potassium ion transport”, “behavior” and “regulation of neuron projection development”, along with the KEGG pathway “Neuroactive ligand-receptor interaction” (Figure 4c, d). Furthermore, the enrichment analysis from DisGeNET indicated that the gene list was primarily enriched for “mood disorders” and also for “movement disorders” (Figure 4e). For the second significant g-LC component in the positive condition, the identified 1,317 genes similarly exhibited significant enrichment in GO biological processes related to synaptic transmission, ion transport, and neuron projection development, and also displayed significant enrichment in diseases linked to depression and motor (Figure S6c, d).

Genes extracted from the main identified g-LC patterns were also enriched for specific cell types. The cell type enrichment analysis indicated that endothelial, microglia, excitatory neurons, inhibitory neurons, and oligodendrocyte precursor cells showed significant enrichment in one of the gene sets. Notably, astrocytes were significantly enriched in two gene sets, covering positive genes in both positive and negative conditions (Figure 4f, details in Figure S7).

### Reproducibility of the SC pattern and transcriptomic profile

We validated the SC pattern associated with physical activity and depressive mood, along with its linked transcriptional profiles, in an independent MDD dataset (MDD-2: N=3,496, MDD/HC=1,775/1,721). The characteristics of the participants in the MDD-2 dataset are presented in Table 1. Using PLS analysis on the MDD-2 dataset, we derived a SC pattern positively correlated with MET score (*r* = 0.52, *p* = 5.8e-246) and showed significant differences between MDD and HC (Cohen’s *d* = -0.07, *p* = 3.4e-2) (Figure 5a). This pattern negatively correlated with depressive mood score (*r* = -0.05, *p* = 1.4e-3) and highlighted the connections between reward-related and motor-related regions as well (Figure 5b). In the genetic scale, the enriched pathways are listed in Figure 5c, which showed overlap with results from the MDD-1 dataset, including “regulation of G protein-coupled receptor signaling pathway”, “behavior”, “regulation of nervous system development”, “response to hormone”, and “head development”. Furthermore, the enrichment analysis of diseases also replicated “mood disorders” as the top-ranking disease (Figure 5d).

**Figure 5.**
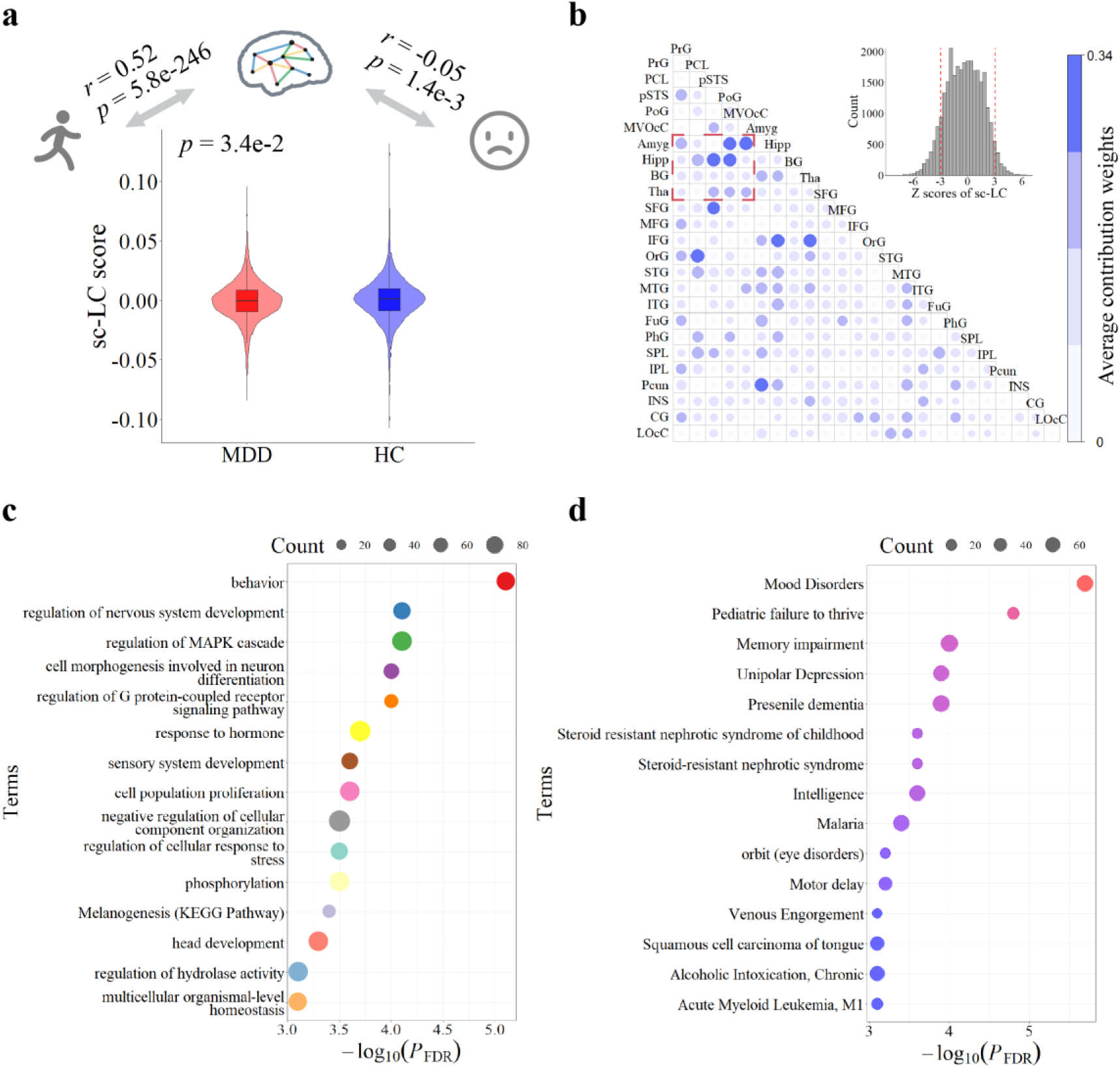
Validation analysis in the MDD-2 dataset. **a**, Reproducible correlation analysis and group difference comparison. The sc-LC score showed a positive correlation with physical activity (*r* = 0.52, *p* = 5.8e-246) and was lower in the MDD group compared to the HC group (*p* = 3.4e-2), with a negative correlation with depressive mood score (*r* = -0.05, *p* = 1.4e-3). **b**, Average contribution weights calculated based on reliable connections (upper right). **c**, The top 15 representative enriched terms of GO biological processes and KEGG pathway for the derived g-LC gene list in the MDD-2 dataset. **d**, The top 15 representative terms enriched in DisGeNET for the derived g-LC gene list in the MDD-2 dataset.

### Generalizability in the BD dataset

We used the identical strategy to assess generalizability in the BD dataset. PLS analysis revealed a SC pattern correlated with physical activity (*r* = 0.89, *p* = 5.6e-28) and showed significant differences between BD and HC (Cohen’s *d* = -0.46, *p* = 4.0e-2) (Figure S8a, b). This pattern similarly highlighted significant contributions between reward-related and motor-related regions (Figure S8c). However, we did not observe a significant association with depressive mood score, which may be attributed to the small sample size (43BDs/38HCs). Furthermore, enrichment analyses demonstrated high concordance with findings from the MDD-1 dataset (see supplemental material and Figure S8 for more details). These results supported potential generalizability to BD.

## Discussion

To our knowledge, this is the first multi-scale multi-level analysis to explore how physical activity is linked to lower human depressive status by combining physical activity score, depressive score, SC and transcriptome data. We established a replicable SC pattern linking motor-related regions and reward-related regions, which was associated with increased physical activity scores and decreased depressive mood scores. The imaging pattern was further correlated with several neurotransmitter receptors, such as serotonin and GABA receptors, and highlighted biological pathways like synaptic signaling and the astrocytes cell type. The imaging and transcriptomic foundations were validated in the MDD-2 dataset and demonstrated generalizability to BD. These findings help better elucidate the imaging mechanism of how physical activity improves depressive mood and the possible underlying biological basis.

At the neuroimaging level, our study revealed a structural interaction between motor-related and reward-related networks. The motor-related regions mainly involved the precentral gyrus, paracentral lobule, postcentral gyrus, posterior superior temporal sulcus, and medioventral occipital cortex, with existing research showing increased volumes in these regions after physical activity^63,64^. The regions implicated in reward processing encompassed the amygdala, hippocampus, basal ganglia, and thalamus, the impairments of which were also observed in individuals with MDD^9,10,65^. Based on the small sample of longitudinal data involving exercise interventions on healthy sedentary subjects, a study focusing on the FC between amygdala and whole brain revealed that changes in amygdala-temporal pole FC were driven by exercise-induced changes in fitness^66^. Another longitudinal study found that hippocampus-precentral gyrus and somatosensory network-thalamus FC were closely associated with changes in aerobic capacity^67^. These two longitudinal analyses partially corroborate our findings regarding the interaction between the motor and reward networks. Despite significant correlations among physical activity, SC pattern and depressive mood, mediation analysis did not reveal a causal relationship. This lack of causality may be due to the cross-sectional nature of our data rather than a longitudinal approach and the effect of exercise on mood may result from multiple neurobiological pathways working together^68^, which could partly explain the difficulty in capturing this effect in a single mediation model.

The association analysis revealed a significant spatial correlation between the identified imaging pattern and the distributions of several neurotransmitter receptors, with serotonin and GABA receptors showing the strongest correlations. Agonists targeting 5-HT1a and 5-HT2a receptors have been reported to promote dopamine release, which could potentially contribute to the antidepressant effects^69^. Indeed, 5-HT1a receptors have already served as a therapeutic target using drugs such as buspirone^70^. Furthermore, 5-HT2a receptors have been found to mediate the neuroplasticity-promoting and antidepressant-like effects of psychedelics^71^, while GABAa^72^ receptors have also shown antidepressant potential. These studies suggest that the identified neurotransmitter receptors may serve as targets for antidepressant therapies, which might contribute to the link between physical activity and depressive mood.

Our investigation delineated the multifaceted mechanisms involved in the connection between exercise and depressive mood, encompassing not only structural interactions and neurotransmitter receptors, but also further pathway and cell type enrichment underpinnings. The identified pathways were primarily involved in synaptic signaling and neuron development, demonstrating reproducibility in the MDD-1 and MDD-2 datasets. Aberrant synaptic signaling can perturb the release, reception, and reuptake of neurotransmitters, such as serotonin, dopamine, GABA, and glutamate, which are closely associated with depression^73,74^. Previous research indicated that treadmill exercise improved inhibitory synaptic transmission in the prefrontal cortex of mice^75^. Furthermore, depression has been linked to abnormal neuron development^76^, while exercise has been shown to positively influence this process^77^. The cell types identified in the enrichment analysis primarily involved astrocytes. Astrocytes play a crucial role in regulating synaptic neurotransmission and promoting synaptic development^78^. In depression, a decrease in the number and volume of astrocytes have been observed in cortical and hippocampal regions^79,80^. Interestingly, physical activity and antidepressant medications, such as fluoxetine and ketamine, have been found to alleviate depressive symptoms, potentially through the protection of astrocytes^81,82^. Thus, astrocytes may be a potential target for therapeutic strategies, including physical activity and antidepressant medications. In summary, our findings on pathways and cell types established important biological connections between exercise and depression and highlighted potential therapeutic avenues.

Our reproducibility and generalizability analysis indicated that the identified neural mechanism and genetic underpinnings were verifiable on the independent MDD-2 dataset and could be generalized to BD. Extensive meta-analysis and prospective studies have consistently demonstrated that physical activity interventions effectively alleviate depressive symptoms in BD, but not manic symptoms^83,84^. The similar symptomatology during depressive phases and impairment patterns in individuals with MDD and BD indicate a potential shared neural basis^85^. Our findings support the potential of exercise as an adjunctive intervention to improve depressive symptoms in both MDD and BD, while the stable, generalizable neuroimaging interaction and pathways like synaptic signaling and neuron development provide valuable insights into the shared imaging and biological basis.

Several potential limitations should also be considered for the current study. First, our current investigation was about cross-sectional analysis instead of longitudinal treatment analysis. It primarily yields association findings. Large samples of longitudinal data will help better explore causality for future investigations. Second, the sample size of the BD dataset used in our generalizability analysis was limited. Although we observed a similar or generalizable neural basis between MDD and BD, future research with larger sample sizes of BD subjects is still needed to better elucidate the neural and genetic basis linking physical activity to depressive mood in BD. Third, the UK Biobank cohort predominantly consists of individuals aged 40-69, with a large proportion of white individuals. To assess the broader applicability of these findings, further replication studies are needed in younger cohorts and among multi-ethnic populations.

## Conclusions

This study integrated multi-level transcriptome-brain imaging-behavior to investigate the neuroimaging basis and biological mechanism underlying the effects of physical activity on human depressive mood. Our findings emphasized the pivotal role played by the interaction between the motor-related and reward-related networks in elucidating the correlation between exercise and depressive mood. The interaction was further linked with serotonin and GABA receptors, and regulated through synaptic signaling and the astrocytes cell type. These findings engender a comprehensive comprehension of the multifaceted mechanisms behind the association, and concurrently furnish potential targets for therapeutic interventions of depression.

## Data availability

The UK Biobank data are accessible to researchers by submitting a data request proposal (https://www.ukbiobank.ac.uk/). Further details on registration for data access are available at http://www.ukbiobank.ac.uk/register-apply/. The Brainnetome atlas is publicly available via https://atlas.brainnetome.org/download.html. Neurotransmitter receptor maps can be downloaded from the neuromaps toolbox (https://netneurolab.github.io/neuromaps/). Gene- expression data were sourced from the Allen Human Brain Atlas, a publicly accessible resource (https://human.brain-map.org/static/download). Cell-specific gene set lists can be downloaded from https://github.com/jms290/PolySyn_MSNs/blob/master/Data/AHBA/celltypes_PSP.csv.

## Code availability

The dMRI data were processed using FSL (https://fsl.fmrib.ox.ac.uk/fsl/fslwiki). The code for PLS analysis is available at https://github.com/SarahMorgan/Morphometric_Similarity_SZ. The processing and analysis of neurotransmitter receptor maps can be found at https://github.com/netneurolab/hansen_receptors and https://netneurolab.github.io/neuromaps/. Preprocessing of gene expression data was conducted using the abagen toolbox (https://github.com/rmarkello/abagen). Gene enrichment analysis was conducted using Metascape (https://metascape.org/gp/index.html#/main/step1).

## Acknowledgements

This work was supported by STI2030-Major Projects 2021ZD0200200; National Natural Science Foundation of China (grant nos. 82001450, 82151307); China Postdoctoral Science Foundation (BX20200364, GZC20232999 and 2024M753502); the Chinese Academy of Sciences, Science and Technology Service Network Initiative (KFJ-STS-ZDTP-078); the Strategic Priority Research Program of the Chinese Academy of Sciences (XDB32030200).

