## supplemental material for "Physical Activity and Depressive Mood Shared the Structural Connectivity between Motor and Reward Network: a population-based study from the UK Biobank"

### Supplementary Methods

#### Participants

The UK Biobank recruited over 500,000 participants from 22 centers across the United Kingdom between 2006 and 2010, and collected a variety of phenotypic and health-related information[1]. Since 2014, additional collection of imaging data and further assessment of behavioral information were conducted on a subset of recruited subjects. Ethical approval was granted by the North West Multi-centre Research Ethics Committee and written informed consent was obtained from all participants.

According to the International Classification of Disease version 10 (ICD-10) codes from hospital inpatient records, 18,429 participants diagnosed with major depression disorder (MDD) (ICD-10: F32/F33) and without a concurrent diagnosis of bipolar disorder (ICD-10: F31) were selected from all individuals. After excluding participants missing physical activity data (N=4,536), depression measurement data (N=2,521) and/or covariate variables (N=3,703), 10,718 participants were remained for association analysis.

In the UK Biobank cohort of all participants, diffusion imaging data were collected from 39,015 subjects. From this group, 23,135 individuals were identified as healthy controls (HC) with no reported records of mental disorders or diseases of the nervous system according to ICD-10 (Field 130836-131127). Furthermore, within the group of participants with diffusion imaging data, 783 participants were also in the cohort of MDD based on the records of hospital inpatient. Subjects with unusable diffusion MRI data (N=40) were excluded from the study, resulting in a group of 743 individuals diagnosed with MDD. HCs were selected from the healthy control (HC) cohort based on the matching criteria including age, sex, and site. Certain participants were excluded from both the MDD and HC groups due to missing physical activity data (N=218), depression measurement data (N=116), covariate variables (N=199), and/or outlier data (N=34). Following this selection process, the MDD-1 dataset consisted of 1,027 participants (MDD/HC=492/535), which was subsequently utilized for the primary analysis.

#### Transcriptome dataset preprocessing

The microarray data were preprocessed utilizing the toolbox abagen[2]. Firstly, we employed the probe-to-gene reannotation using data from Arnatkeviciute et al.[3]. Probes lacking a valid Entrez ID were excluded from further analysis. Following this, we performed a filtering step to eliminate probes with expression below background in 50% of all samples across donors[4]. For genes that corresponded to multiple probes, representative probes exhibiting the highest correlation of expression values across donors were chosen[5]. The probes selection process above resulted in 15,605 unique probes, each corresponding to a distinct gene. Subsequently, the samples were matched to 246 regions as defined in the Brainnetome Atlas. Any samples that failed to match with a specific brain region were removed. To reduce the risk of inaccurate assignment, the matching process was restricted according to both hemisphere and gross structural divisions. Gene expression values were normalized across genes using scaled robust sigmoid. Identical normalization was further performed across the samples within cortex, subcortex, and cerebellum separately[6]. Finally, we combined the samples within each brain region by computing the average expression data across all samples belonging to that region for each donor, followed by averaging across donors. Since the absence of tissue samples in 11 out of the 246 brain regions, a gene expression matrix of size 235×15,605 was obtained.

#### Partial least squares analysis

Partial least squares (PLS) regression is a multiple linear regression technique that not only considers the correlation between the principal components of both predictor (X) and response (Y) variables, but also accounts for their individual variances. This approach thereby mitigates the challenge of multicollinearity in high-dimensional data. In the present study, PLS regression was employed to determine the relationship linking structural connectivity (SC) with physical activity, as well as the association between gene expression and the identified SC pattern.

In the association analysis between SC and physical activity, the predictor variables comprised the whole-brain structural connections, denoted as X (participants × SC values), while the response variables consisted of the total MET scores, denoted as Y (participants × 1). Subsequently, in the neuroimaging-transcriptome association analysis, the gene expression data were stored in X (regions × gene expression) and the features of the SC pattern were stored in Y (regions × 1). Both X and Y matrices were normalized by transforming them into respective Z-scores denoted by X0 and Y0.

The PLS analysis procedure is as follows[7]. The covariance matrix between the normalized predictor variables ($X_0$) and the normalized response variables ($Y_0$) is calculated as:

$$R_1 = X_0^T Y_0$$

The singular value decomposition (SVD) is computed on the covariance matrix $R_1$:

$$R_1 = W_1 \Delta_1 C_1^T$$

This yields the eigenvectors $W_1$ and $C_1$, where the first pair is denoted as $w_1$ and $c_1$. $\Delta_1$ is the diagonal matrix containing the eigenvalues. The first latent component (LC) scores of X and Y are calculated by:

$$t_1 = X_0 w_1$$

$$u_1 = Y_0 c_1$$

The predictor loadings are calculated by:

$$p_1 = X_0^T t_1$$

Thus, we can reconstruct $X_0$ using $t_1 p_1^T$. The reconstruction of $Y_0$ based on the LC of X is obtained by:

$$\widehat{Y_1} = u_1 c_1^T = t_1 \mathrm{b}_1 c_1^T \quad with \quad \mathrm{b}_1 = t_1^T u_1$$

The residuals after reconstructing are used as $X_1$ and $Y_1$, which play the role in $X_0$ and $Y_0$ for the next iteration. The iteration continues until a specified number of components or the rank of X is reached. The vectors $t_i, u_i, p_i, w_i$, and $c_i$, obtained at each iteration, are stored in their respective matrices T, U, P, W, and C. Matrices T and U contain the LC scores of X and Y, respectively. Predictor loadings are represented in matrix P, while matrices W and C capture the LC weights for X and Y, respectively.

To test the statistical significance of the LCs, we performed 1000 permutation tests. In each permutation test, we randomly shuffled Y and repeated the entire procedure of PLS analysis. In the SC-behavior association analysis, we conducted the permutations separately within the two groups (the MDD group and the HC group) due to the significant differences in physical activity observed between these groups[8]. Furthermore, bootstrap was performed 1000 times to estimate the variability of the LC weights, which was represented using Z score, calculated by dividing the PLS weight of each feature in X by its bootstrapped standard error.

#### Antidepressant usage

To assess the impact of medication on our findings, we divided participants into medicated and unmedicated groups based on antidepressant usage. The UK Biobank provides verbal interview data on prescription medication use (Field 20003). Utilizing Data-Coding 4 from Field 20003 and following Glanville et al.[9], we defined "Antidepressant Usage" using the antidepressant codes: 1140879616, 1140921600, 1140879540, 1140867878, 1140916282, 1140909806, 1140867888, 1141152732, 1141180212, 1140879634, 1140867876, 1140882236, 1141190158, 1141200564, 1140867726, 1140879620, 1140867818, 1140879630, 1140879628, 1141151946, 1140867948, 1140867624, 1140867756, 1140867884, 1141151978, 1141152736, 1141201834, 1140867690, 1140867640, 1140867920, 1140867850, 1140879544, 1141200570,

1140867934, 1140867758, 1140867914, 1140867820, 1141151982, 1140882244, 1140879556, 1140867852, 1140867860, 1140917460, 1140867938, 1140867856, 1140867922, 1140910820, 1140882312, 1140867944, 1140867784, 1140867812, 1140867668. Participants who reported taking any of the mentioned medications were classified as the medicated group. The unmedicated group was consisted of participants who had not used these medications. A total of 238 medicated and 254 unmedicated participants were included in the subsequent analysis. Detailed demographic information is presented in Table S3.

#### Validation and generalization analysis

PLS regression was performed separately on the MDD-2 and BD datasets to evaluate the reproducibility and generalizability of the findings, with total MET scores serving as the response variables and inter-regional SC features as the predictor variables. Using the same strategy as in the MDD-1 dataset, we derived the sc-LC from PLS and utilized the sc-LC score for group difference comparison and association analysis with depressive mood score. Furthermore, in the genetic analysis, we applied the same strategy to obtain the g-LC, which exhibited statistical significance and the largest explained variance. Genes with $|Z| > 3$ after bootstrapping were retained and gene enrichment analysis was performed using Metascape. FDR correction was used for multiple comparison corrections.

### Supplementary Results

#### Association of sc-LC score with various intensities of physical activity

According to the International Physical Activity Questionnaire (IPAQ) short form[10], total physical activity consists of walking, moderate activity, and vigorous activity. We examined the correlation between the sc-LC score and physical activity across these three specific activity types. Our findings revealed that higher sc-LC score was significantly correlated with increased MET score in walking ($r$ = 0.48, $p$ = 1.5e-60), moderate activity ($r$ = 0.55, $p$ = 1.4e-81), and vigorous activity ($r$ = 0.45, $p$ = 1.7e-51) (Figure S2). Among these different intensities of exercise, moderate activity exhibited the strongest correlation with sc-LC score, suggesting that engaging in moderate activity may have a greater potential for improving depressive mood.

#### Mediation Analysis

Given that the sc-LC score was simultaneously associated with the MET score and the depressive mood score, a mediation analysis was performed to explore whether a mediation effect of $MET \xrightarrow{sc-LC} mood$ may occur. The PROCESS macro v3 for SPSS was used for mediation analyses[11,12]. Specifically, MET score served as the independent variable, the sc-LC score as the mediator, and depressive mood score as the dependent variable. The indirect effect (CI = [-0.01, 0.09]) of the MET score on the depressive mood score via the sc-LC score was not significant. In the regression model examining both physical activity and the sc-LC score on depressive mood, the effect of the sc-LC score on depressive mood did not reach significant (CI = [-1.64, 11.35]). This could potentially be influenced by the cross-sectional nature of the data rather than being longitudinal.

#### Separate PLS analyses within MDD and HC group

Separate PLS analyses were conducted for MDD/HC separately. Figure S3a indicates a PLS component showing significant correlation with the MET score ($r$ = 0.68, $p$ = 6.0e-69) identified on MDD only. The weights of the component exhibited a high correlation with the original pattern ($r$ = 0.69, $p$ < 0.001). Figure S3b indicates a PLS component showing significant correlation with the MET score ($r$ = 0.61, $p$ = 4.6e-55) identified on HC only. The weights also exhibited a high correlation with the original pattern ($r$ = 0.67, $p$ < 0.001). Both populations similarly highlighted the connections between the motor and reward network.

These results indicate that separate PLS analyses yielded similar findings.

#### Separate PLS analyses within medicated and unmedicated groups

To assess the impact of medication on our findings, separate PLS analyses were performed for medicated and unmedicated groups. In the medicated group, we identified a PLS component showing significant correlation with physical activity ($r = 0.61$, $p = 1.2e\text{-}25$). Similarly, key connections were observed between reward-related and motor-related regions (Fig. S5a). The component weights showed a significant correlation with the originally identified pattern ($r = 0.40$, $p < 0.001$). In the unmedicated group, a PLS component showing significant correlation with the MET score ($r = 0.62$, $p = 2.8e\text{-}28$) was identified, with contributing structural connections primarily between reward-related and motor-related regions (Fig. S5b). The component weights similarly exhibited a significant correlation with the original pattern ($r = 0.43$, $p < 0.001$). These results suggest that the influence of medication treatment on the outcomes was relatively minor, as the findings were highly consistent across both medicated and unmedicated groups.

#### Validation analyses of gene expression results

Since the limited inclusion of right hemispherical data in the AHBA dataset (only 2 donors), potential bias was a concern. To address this, we added a mirroring step for microarray expression samples across hemispheres during the AHBA dataset preprocessing[2]. PLS regression was employed on the new transcriptome matrix. Subsequently, we computed the correlation between the weights of the gene component before and after the mirroring process. The results demonstrated a high correlation with our main results in both positive and negative conditions ($r = 0.80$, $p < 0.001$ for positive condition; $r = 0.75$, $p < 0.001$ for negative condition).

Furthermore, among the 6 donors in the AHBA dataset, 5 were male, and 1 was female. To investigate the potential impact of sex on the results, we specifically selected the 5 male donors for analysis. Similar to the hemisphere validation, we calculated the correlation between the weights of the gene component before and after this selection. The results still revealed a high correlation with our main results in both positive and negative conditions ($r = 0.96$, $p < 0.001$ for positive condition; $r = 0.95$, $p < 0.001$ for negative condition). Collectively, our validation results indicate that the findings based on the AHBA dataset were stable and robust to hemisphere and sex.

#### Reproducibility and generalizability of transcriptomic profile

In the MDD-2 dataset, the identified component (g-LC) which explained the largest proportion of variance (16%, permuted $p$ = 1.0e-3) and presented the highest correlation with the corresponding pattern ($r$ = 0.40, $p$ = 1.9e-10) was adopted for subsequent enrichment analysis. The pathways enriched by significant contributing genes are listed in Figure 5c, which showed overlap with results from the MDD-1 dataset.

In the BD dataset, a significant g-LC that explained the largest proportion of variance (16%, permuted $p$ = 1.5e-2) and showed the highest correlation with the corresponding SC pattern ($r$ = 0.40, $p$ = 1.3e-10) was identified. The gene list with significant contributions enriched pathways such as “modulation of chemical synaptic transmission”, “synaptic signaling”, “head development”, “neuron projection development”, “regulation of system process”, and “cellular response to hormone stimulus”, demonstrating high concordance with the findings in the MDD-1 dataset (Figure S8d).

### Supplementary Tables

**Table S1. All variables and their fields from UK Biobank used in the current study.**

| Description | Field Id |
|---|---|
| Participants diagnosed with MDD | |
| Date F32 first reported | 130894 |
| Source of report of F32 | 130895 |
| Date F33 first reported | 130896 |
| Source of report of F33 | 130897 |
| Major depression status | 20126 |
| Participants diagnosed with BD | |
| Date F31 first reported | 130892 |
| Source of report of F31 | 130893 |
| Physical activity | |
| Number of days/week of walking | 864 |
| Duration of walking | 874 |
| Number of days/week of moderate activity | 884 |
| Duration of moderate activity | 894 |
| Number of days/week of vigorous activity | 904 |
| Duration of vigorous activity | 914 |
| Depression measurement | |
| Frequency of depressed mood in last 2 weeks | 2050 |
| Frequency of disinterest in last 2 weeks | 2060 |
| Frequency of tenseness in last 2 weeks | 2070 |
| Frequency of tiredness in last 2 weeks | 2080 |
| Age at recruitment | 21022 |
| Year of birth | 34 |
| Month of birth | 52 |
| Date of imaging visit | 53 |
| Sex | 31 |
| Imaging site | 54 |
| Education | 6138 |
| Ethnicity | 21000 |
| Townsend deprivation index | 189 |
| Employment status | 6142<br>20119 |
| Body mass index | 21001 |
| Average total household income before tax | 738 |

Note: Variables at instance 0 (initial assessment visit) were obtained to conduct an association analysis without considering MRI data. Variables at instance 2 (imaging visit) were extracted to perform analyses combining both the variables and imaging data. MDD, major depressive disorder; BD, bipolar disorder.

**Table S2. The detailed information of the neurotransmitter receptor map.**

| | Description | N | Reference(s) |
|---|---|---|---|
| 5-HT1a | way100635 | 35 | Savli et al., 2012[13] |
| 5-HT1b | p943 | 23<br>23 | Gallezot et al., 2010[14]<br>Savli et al., 2012[13] |
| 5-HT2a | altanserin | 19 | Savli et al., 2012[13] |
| 5-HT4 | sb207145 | 59 | Beliveau et al., 2017[15] |
| 5-HT6 | gsk215083 | 30 | Radhakrishnan et al., 2018[16] |
| D1 | sch23390 | 13 | Kaller et al., 2017[17] |
| D2 | flb457 | 55<br>37 | Sandiego et al., 2015[18]<br>Smith et al., 2017[19] |
| α4β2 | flubatine | 30 | Hillmer et al., 2016[20] |
| M1 | lsn3172176 | 24 | Naganawa et al., 2020[21] |
| GABAa | flumazenil | 6 | Dukart et al., 2018[22] |
| mGluR5 | abp688 | 28<br>22<br>73 | Dubois et al., 2016[23]<br>Rosaneto[24]<br>Smart et al., 2019[25] |
| CB1 | omar | 77 | Normandin et al., 2015[26] |
| MOR | carfentanil | 204<br>39 | Kantonen et al., 2020[27]<br>Turtonen et al., 2021[28] |
| H3 | gsk189254 | 8 | Gallezot et al., 2017[29] |

**Table S3. Demographic characteristics of participants in the medicated and unmedicated groups.**

| | MDD in MDD-1 Dataset | | |
|---|---|---|---|
| | **Medicated** | **Unmedicated** | **p** |
| **Subjects** | **238** | **254** | |
| **Age, Years** | 61.49 ± 7.55 | 62.67 ± 7.74 | 0.09 |
| **Sex (Male/Female)** | 34%/66% | 43%/57% | 0.05 |
| **Education attainment (High/Low)** | 47%/53% | 48%/52% | 0.90 |
| **Ethnicity (White/Non-white)** | 99%/1% | 98%/2% | 0.53 |
| **Townsend deprivation index** | -1.62 ± 2.81 | -1.12 ± 3.14 | 0.06 |
| **Employment status (In paid employment/Other)** | 37%/63% | 40%/60% | 0.47 |
| **Body mass index** | 28.82 ± 5.30 | 28.09 ± 5.11 | 0.12 |
| **Average total household income before tax** | | | |
| **Less than 18,000** | 23% | 24% | 0.34 |
| **18,000 to 30,999** | 31% | 33% | |
| **31,000 to 51,999** | 29% | 23% | |
| **52,000 to 100,000** | 13% | 17% | |
| **Greater than 100,000** | 4% | 3% | |
| **Physical activity, MET-mins/week** | 1928 ± 2034 | 2468 ± 2277 | <0.01 |
| **Depressive mood score** | 3.96 ± 3.47 | 2.79 ± 2.97 | <0.001 |

**Table S4. The explained variance and statistical significance of the first five PLS components in the positive condition and negative condition.**

| | | | | | | |
|---|---|---|---|---|---|---|
| Positive condition | PLS component | 1 | 2 | 3 | 4 | 5 |
| | The percentage of variance explained by PLS component | 13% | 16% | 9% | 11% | 7% |
| | permuted *p* | 4.3e-2 | 4.4e-2 | 0.63 | 0.53 | 0.80 |
| Negative condition | PLS component | 1 | 2 | 3 | 4 | 5 |
| | The percentage of variance explained by PLS component | 17% | 13% | 8% | 10% | 9% |
| | permuted *p* | 2.0e-3 | 0.25 | 0.81 | 0.69 | 0.53 |

### Supplementary Figures

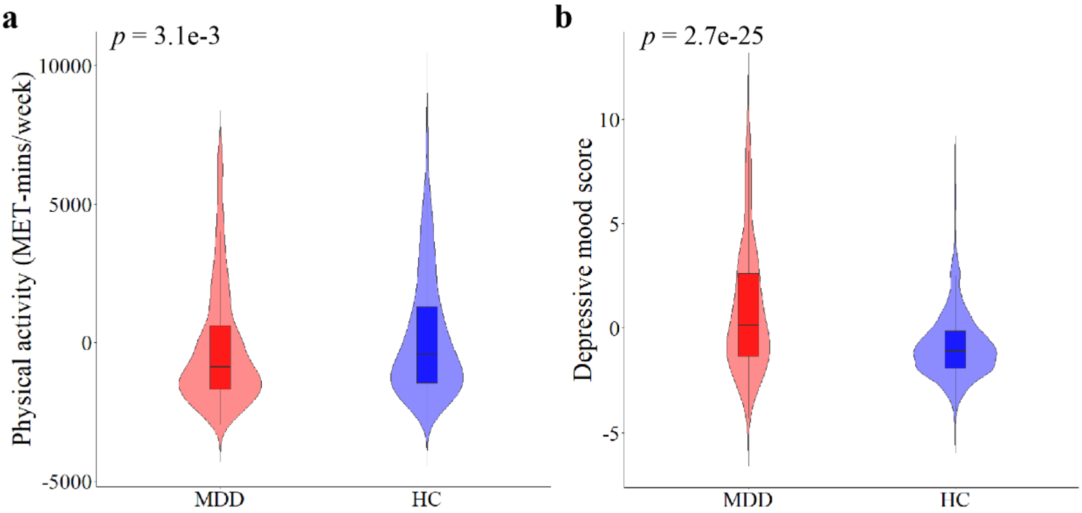


**Figure S1.** Two-sample t-test of **a**, physical activity and **b**, depressive mood score between MDDs and HCs based on the MDD-1 dataset.

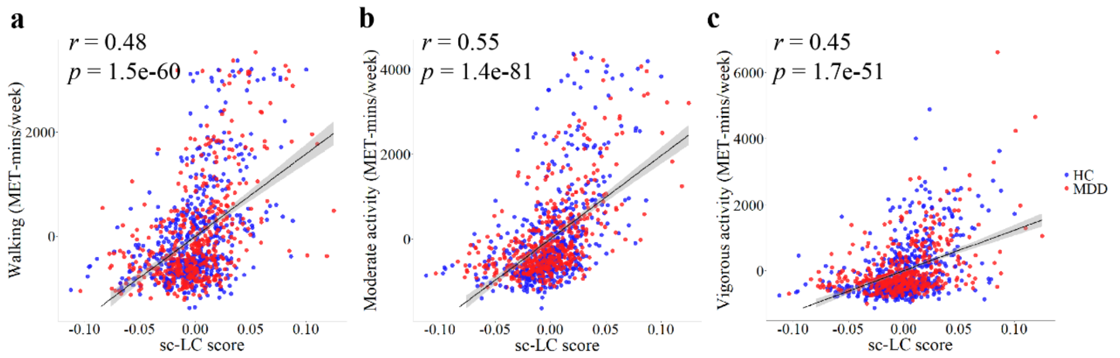


**Figure S2. Association between sc-LC score and various intensities of physical activity (MET score). a**, Association of sc-LC score with walking. **b**, Association of sc-LC score with moderate activity. **c**, Association of sc-LC score with vigorous activity. Moderate activities encompass physical endeavors such as cycling at normal pace, transporting light loads. Vigorous activities are characterized by their ability to induce perspiration and heavy breathing, exemplified by activities such as fast cycling, participating in aerobics, and heavy lifting.

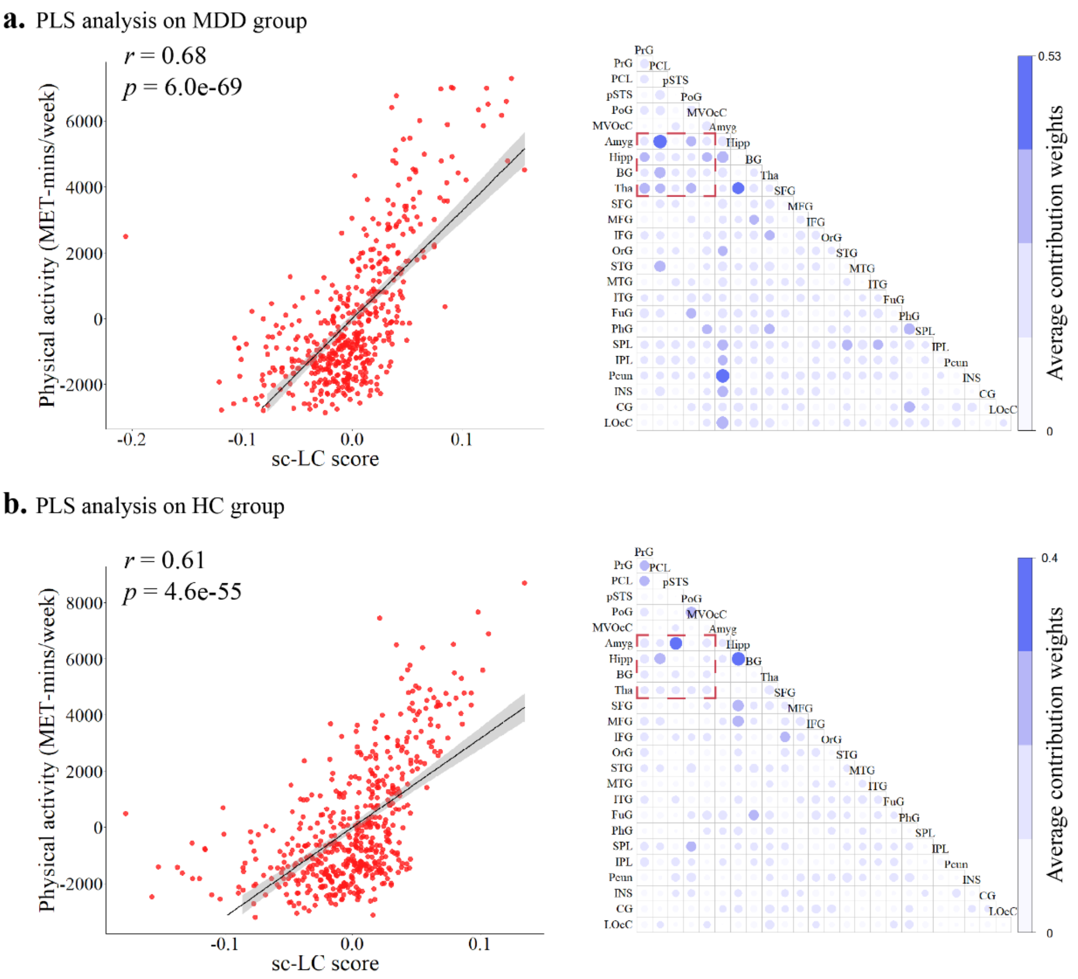


**Figure S3. Conducting separate PLS analyses within MDD and HC groups. a,** PLS analysis on MDD group. **b,** PLS analysis on HC group.

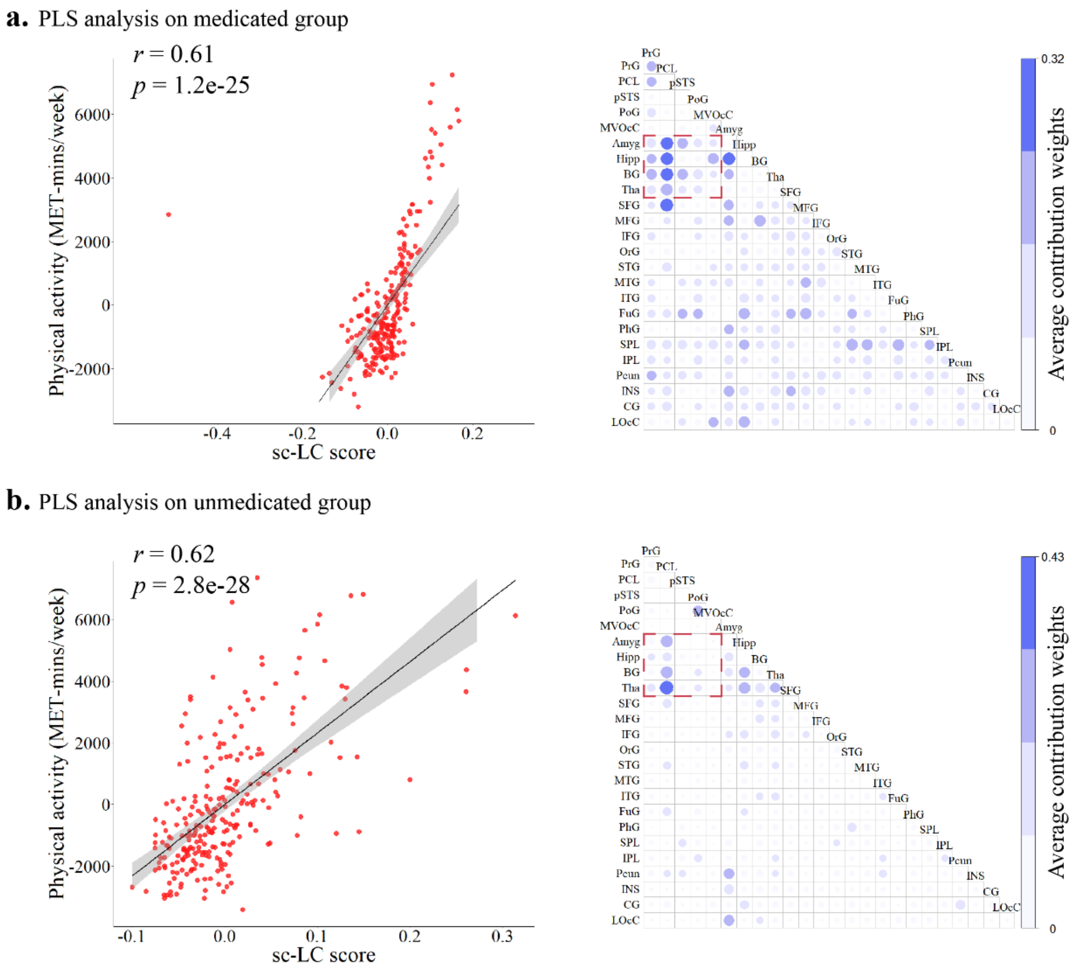


**Figure S4. Conducting separate PLS analyses within medicated and unmedicated groups. a,** PLS analysis on medicated group. **b,** PLS analysis on unmedicated group.

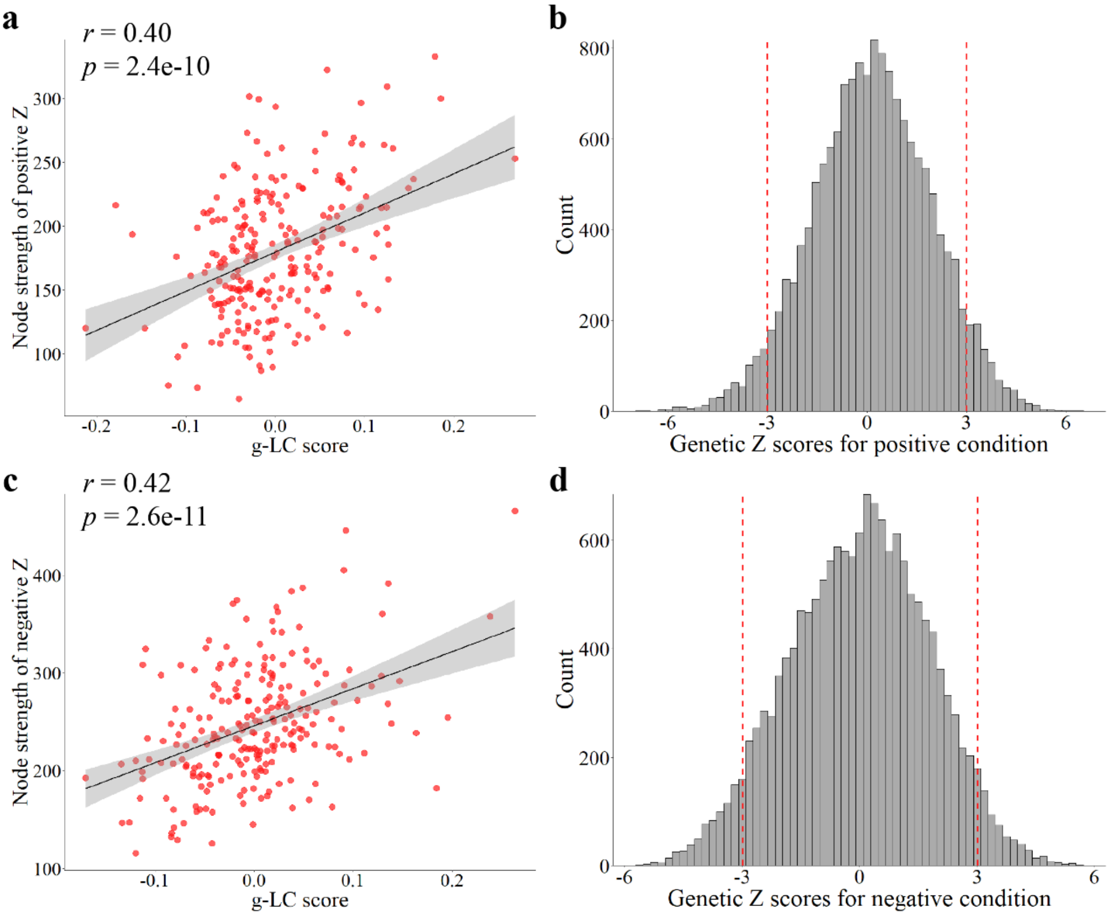


**Figure S5. Association between g-LC score and the sc-LC pattern. a**, Association between g-LC score (positive condition) and node strength of positive Z scores for sc-LC. The g-LC score was calculated by multiplying the weights of the g-LC by 15,605 gene expression scores for each brain region. **b**, The distribution of genetic Z scores for positive condition. Bootstrapping was conducted 1,000 times to estimate genetic Z scores. Among the 15,605 genes, 853 genes exhibited Z scores > 3, and 634 genes displayed Z scores < -3. **c**, Association between g-LC score (negative condition) and node strength of negative Z scores. **d**, The distribution of genetic Z scores for negative condition. Bootstrapping was conducted 1,000 times to estimate genetic Z scores. Among the 15,605 genes, 616 genes exhibited Z scores > 3, and 778 genes displayed Z scores < -3.

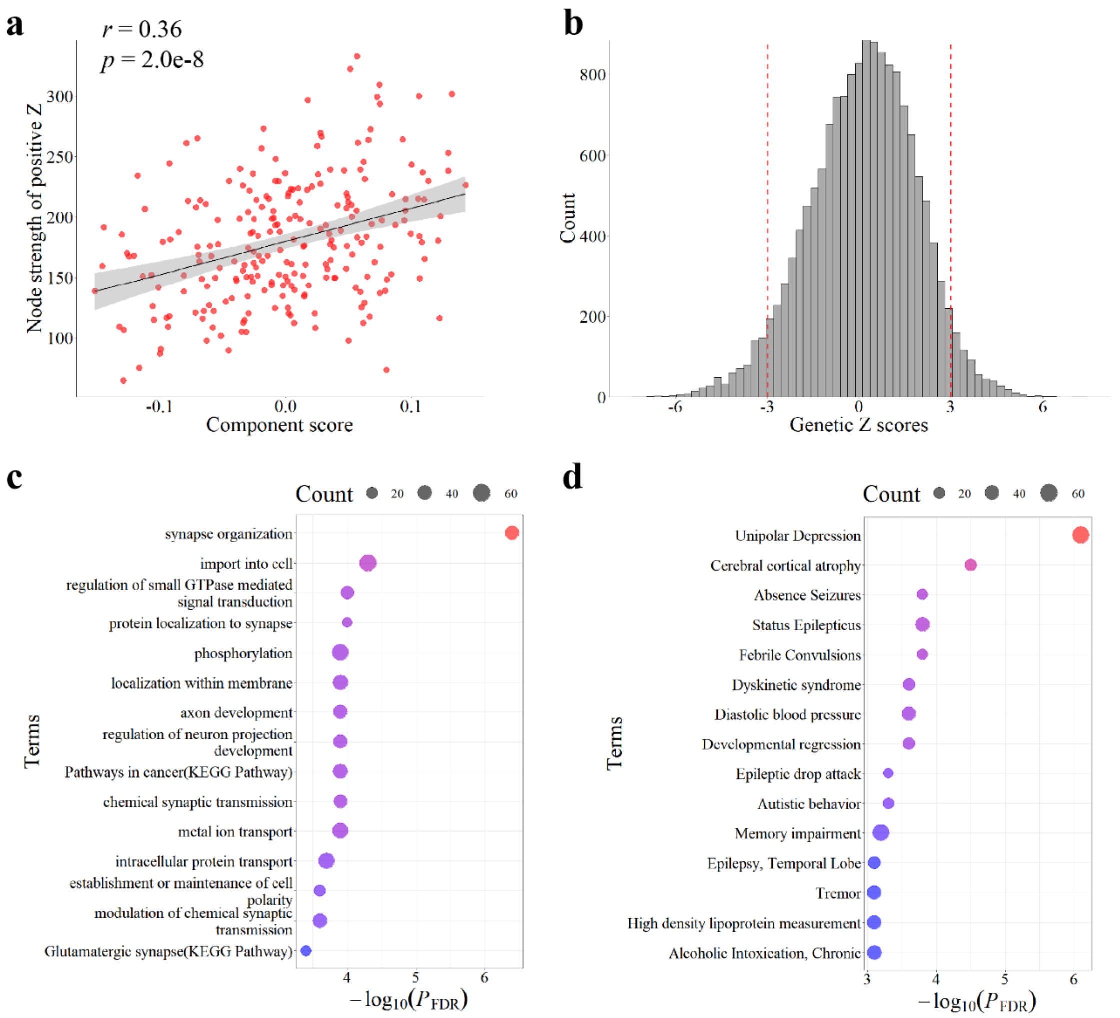


**Figure S6. Enrichment analysis on the gene list for another component in the positive condition. a**, Association between another component and the corresponding pattern in the positive condition. The component score was calculated by multiplying the weights of the component by 15,605 gene expression scores for each brain region. **b**, The distribution of genetic Z scores for this component. **c**, The top 15 representative enriched terms of GO biological processes and KEGG pathway for the gene list. **d**, The top 15 representative terms enriched in DisGeNET for the gene list. The size of each circle corresponds to the number of genes involved in the given ontology terms.

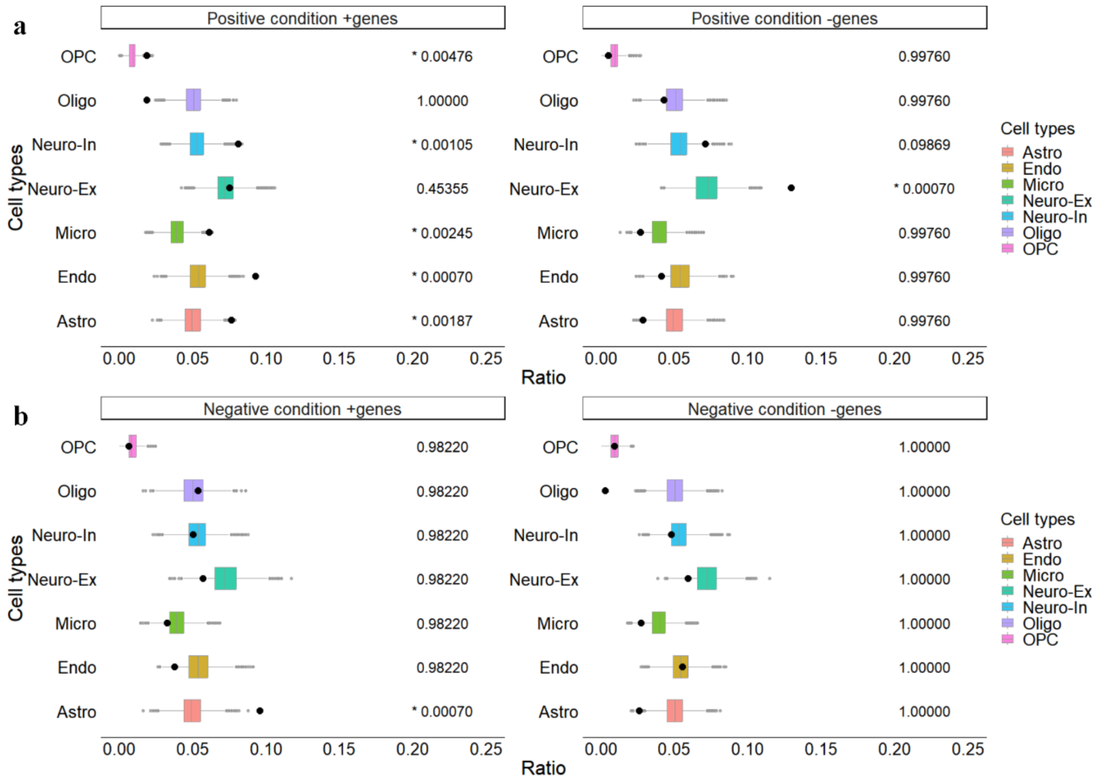


**Figure S7. Cell types analysis. a**, Cell types enrichment for gene lists in positive condition (derived from positive Z scores of sc-LC). **b**, Cell types enrichment for gene lists in negative condition (derived from negative Z scores of sc-LC). For each condition, genes exhibiting a positive correlation with the node strength of Z scores were referred to as "+genes", whereas those demonstrating a negative correlation were termed "-genes". The ratio was determined by dividing the number of genes that overlapped between a specific cell type's gene set and our gene list by the total number of genes included in our gene list. The p-value was derived using a permutation test and subsequently adjusted by multiple comparison using the FDR correction. Astro, astrocytes; Endo, endothelial cells; Micro, microglia; Neuro-Ex, excitatory neurons; Neuro-In, inhibitory neurons; Oligo, oligodendrocytes; OPC, oligodendrocyte precursor cells.

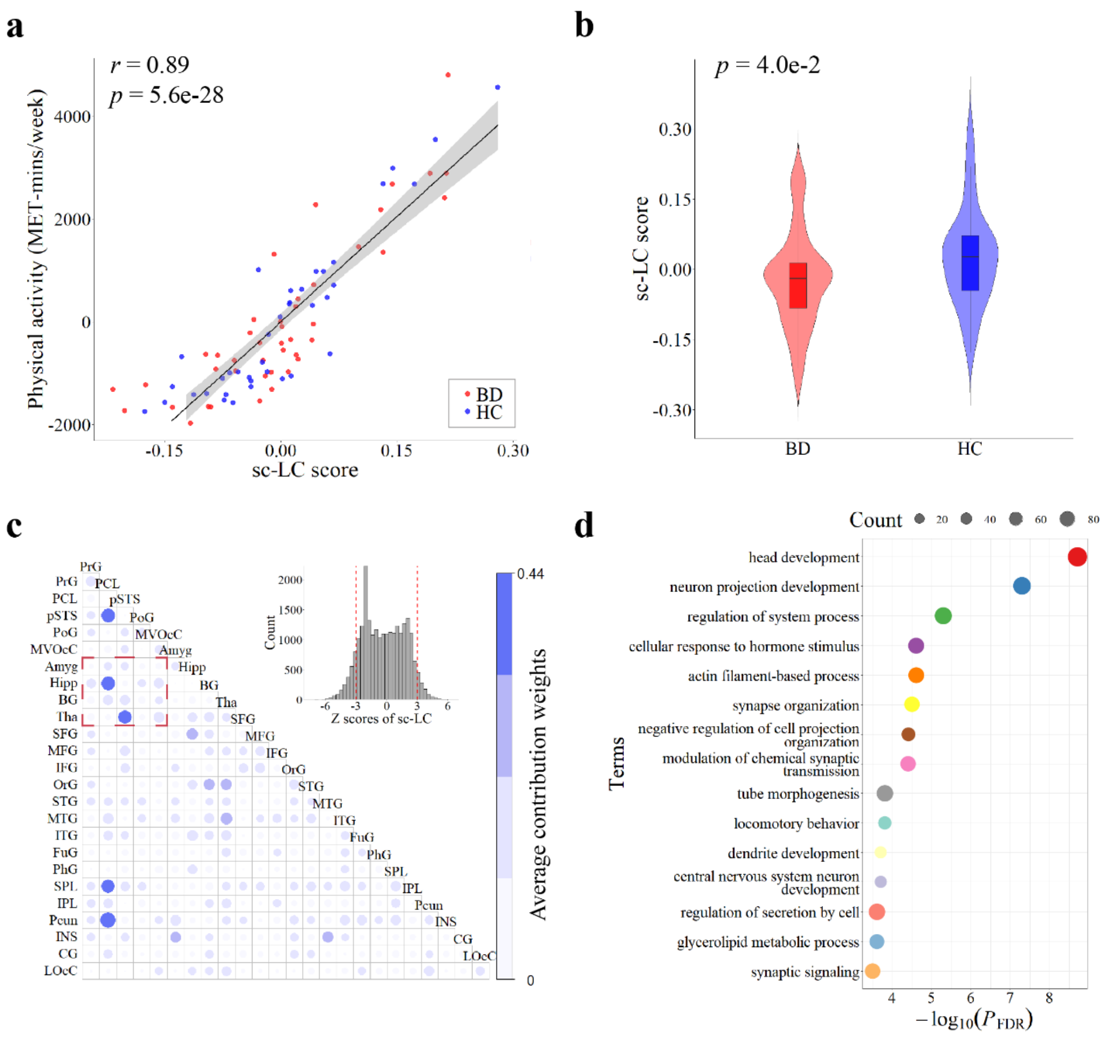


**Figure S8. Generalization analysis in the BD dataset. a**, Correlation between the sc-LC score and physical activity (MET score). **b**, Two-sample two-tailed t-test of sc-LC score between the BDs and the HCs. **c**, Average contribution weights calculated based on reliable connections (upper right). **d**, The top 15 representative enriched terms of GO biological processes and KEGG pathway for the derived g-LC gene list in the BD dataset.

23. DuBois JM, Rousset OG, Rowley J, et al. Characterization of age/sex and the regional distribution of mGluR5 availability in the healthy human brain measured

by high-resolution [11C]ABP688 PET. *Eur J Nucl Med Mol Imaging*. 2016;43(1):152-162. doi:10.1007/s00259-015-3167-6
